# Selenocysteine tRNA methylation promotes oxidative stress resistance in melanoma metastasis

**DOI:** 10.1101/2023.10.02.560568

**Authors:** Leona A. Nease, Kellsey P. Church, Ines Delclaux, Shino Murakami, Maider Astorkia, Marwa Zerhouni, Graciela Cascio, Riley O. Hughes, Kelsey N. Aguirre, Paul Zumbo, Lukas E. Dow, Samie Jaffrey, Doron Betel, Elena Piskounova

## Abstract

Selenocysteine-containing proteins play a central role in redox homeostasis. Their translation is a highly regulated process, dependent upon two tRNA^Sec^ isodecoders differing by a single 2’-O-ribose methylation, called Um34. We characterized FTSJ1 as the Um34 methyltransferase and show that its activity is required for efficient selenocysteine insertion at the UGA stop codon during translation. Specifically, Loss of Um34 leads to ribosomal stalling and decreased UGA recoding. FTSJ1-deficient cells are more sensitive to oxidative stress and have decreased metastatic colonization in xenograft models of melanoma metastasis. We found that FTSJ1 mediates efficient translation of selenoproteins essential for the cellular antioxidant response. Our findings uncover a role for tRNA^Sec^ Um34 modification in oxidative stress resistance and highlight FTSJ1 as a potential therapeutic target specific for metastatic disease.

## Introduction

tRNA modifications coordinate dynamic changes in protein translation, enabling rapid adaptation and survival under various stress conditions^1, 2^. Modifications on the tRNA anticodon loop specifically affect translational efficiency and fidelity^3, 4^. Selenocysteine-containing proteins maintain redox and ER homeostasis and their translation is regulated by a single methyl group at the wobble position (U_34_) of the Sec tRNA^5^. Selenocysteine (Sec), the 21^st^ amino acid, is coded for by the UGA stop codon and synthesis of these proteins is energetically costly. Biosynthesis of Sec occurs on its tRNA rather than as a small molecule, and co-translational incorporation of Sec requires a short hairpin structure in the 3’ UTR. This Selenocysteine Incorporation Sequence (SECIS) is bound by a complex of proteins containing a Sec-specific elongation factor (EEFSEC), SECIS binding protein 2 (SECISBP2) and SECp43 which position the Sec tRNA for loading onto the ribosome and enable readthrough of the UGA codon.^6–9^ (Figure 1A).

**Figure 1:**
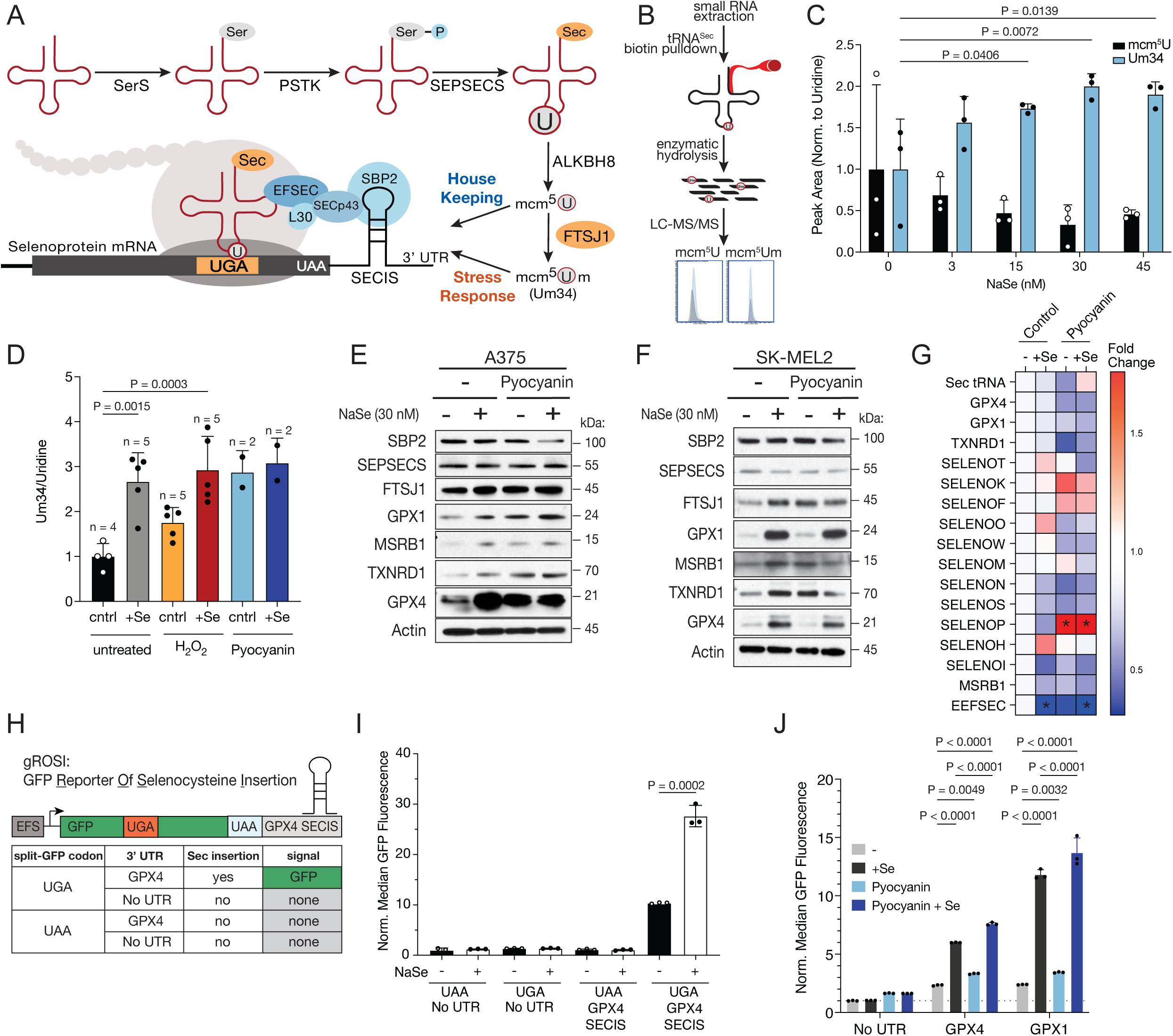
Selenoprotein translation and tRNA^Sec^ Um34 levels increase in response to oxidative stress. (A) Schematic of selenocysteine biosynthesis and selenoprotein translation. Sec is synthesized on its tRNA. The tRNA is then modified at the wobble uridine and the Sec translation complex associated with the SECIS allows for co-translational readthrough of UGA stop codon. (B) Schematic of tRNA^Sec^ isolation and Um34 modification analysis. (C) Quantification of Um34 and mcm^5^U levels in A375 melanoma cells grown in DMEM/F12 medium supplemented with increasing concentrations of selenium measured by LC-MS/MS. n= 3 independent biological experiments. Mean +/- SD. Statistical significance assessed by two-way analysis of variance (ANOVAs) followed by Fisher LSD test for multiple comparisons. (D) Quantification of Um34 modification levels on tRNA^Sec^ in A375 melanoma cells treated with pro-oxidant H_2_O_2_ (100µM) pyocyanin (10µM) in DMEM/F12 media in the presence or absence of 30nM NaSe. n = number of independent samples is indicated in the figure. Mean +/- SD. Statistical significance assessed by one-way ANOVA followed by Fisher LSD test for multiple comparisons. (E) Levels of selenoproteins in A375 melanoma cells and (F) SK-MEL-2 melanoma cells after treatment with NaSe (30nM) and pyocyanin (10uM). Representative from 3 independent experiments shown. (G) Transcript levels of selenoproteins in A375 melanoma cells after treatment with NaSe and pyocyanin measured by qRT-PCR. Mean +/- SD. n=3 independent biological experiments. Statistical significance assessed by two-way analysis of variance (ANOVA) with each condition compared to selenium deficient cells in the DMSO control condition. *p-values: SELENOP Control – Se vs Pyocyanin – Se P=0.0308; SELENOP Control – Se vs Pyocyanin + Se P=0.0210. EEFSEC Control – Se vs Control + Se P= 0.0487; EEFSEC Control – Se vs Pyocyanin + Se P=0.0407. (H) Schematic of the split-seleno-GFP reporter - GFP Reporter Of Selenocysteine Insertion (gROSI). (I) Flow cytometry analysis of split-seleno-GFP reporter (gROSI) expression with different combinations of stop codons UAA or UGA and with No UTR or SECIS from GPX4, in the presence or absence of selenium (30nM NaSe) in A375 melanoma cells. n = 3 independent biological experiments, Mean +/- SD; representative experiment shown; two-sided unpaired t-test, (J) Flow cytometry analysis of GFP fluorescence of gROSI variants without a 3’UTR, with a GPX4 SECIS and with a GPX1 SECIS treated with Pyocyanin (10uM) and NaSe (30nM). n=3 independent biological experiments, Mean +/- SD. Statistical significance measured by two-way ANOVA followed by Fisher LSD test for multiple comparisons.]

Sec tRNA is hypomodified at only 4 positions - pseudoU55 and m^1^A58 are involved in proper folding, while anticodon loop modifications, i^6^A37 and mcm^5^Um34 (Um34), regulate selenoprotein translation^10^. Sec tRNA exists in two forms: with either the precursor mcm^5^U, catalyzed by ALKBH8^11, 12^, or with additional 2’O-ribose methylation generating mcm^5^Um. Um34 is unique to the Sec tRNA and the final step in its maturation, affecting the tertiary structure of the tRNA. Additionally, Um34 is highly responsive to selenium availability, and has been proposed to increase efficiency of Sec insertion^13, 14^. Previous work has suggested these two isodecoders enable the translation of a hierarchy of Sec-containing proteins^15^. This ensures that only essential selenoproteins are synthesized when selenium is limiting, or conversely, non-essential or stress-related selenoproteins are translated when Um34 is increased.

There are 25 known selenoproteins, with 11 involved in the cellular antioxidant response. Glutathione peroxidases (GPX1-4, GPX6) detoxify ROS, with GPX4 specifically inhibiting ferroptosis^16–18^. Thioredoxin reductases (TXNRD1-3) reduce oxidized cysteine residues and MSRB1 reduces oxidized methionine^19, 20,21^. SELENOH is a nuclear oxidoreductase that regulates p53 in response to ROS^22^, and SELENOO is a mitochondrial AMPylase necessary for cellular oxidative stress response^23^. Other selenoproteins maintain ER homeostasis (SELENOF, K, M, N, S and T), regulating and aiding in protein folding^24^. Enzymes DIO1-3 metabolize thyroid hormone^25^ and SELENOP distributes selenium throughout the body^26^. Due to the low pKa of Sec, these enzymes are highly reactive and resist permanent oxidative damage^27, 28^.

We have investigated Um34 function in the context of oxidative stress, given the central role for selenoproteins in oxidative stress resistance. We found Um34 increases with oxidative stress, correlating with increased selenoprotein translation and readthrough of UGA. We characterized FTSJ1 as the Um34 methyltransferase, identifying it as a component of the selenocysteine translation machinery and required for efficient Sec insertion. We found its loss affects translation of antioxidant selenoproteins and sensitizes melanoma cells to oxidative stress. A defining feature of metastasizing cancer cells are high levels of oxidative stress and increased dependence on antioxidant pathways^29^. Our data show that loss of FTSJ1 in melanoma cells has no impact on primary tumor growth but decreases metastatic colonization in distant organs. Our work identifies oxidative stress as a specific context for Um34-mediated selenoprotein translation and highlights FTSJ1 as a potential therapeutic target specific to metastatic disease.

## RESULTS

### Selenoprotein translation and Um34 increase under oxidative stress

Um34 levels were measured by isolating Sec tRNA using a biotinylated oligo probe coupled with LC-MS/MS analysis (Figure 1B and Extended Figure 1A). To establish the effect of extracellular selenium in melanoma cells, A375 cells were depleted of selenium in serum-free DMEM/F12 media. Um34 levels increased with increasing extracellular selenium, while mcm^5^U precursor decreased, showing that selenium regulates Um34 biosynthesis and shifts the ratio between the two Sec tRNA isodecoders in melanoma cells (Figure 1C). Selenoproteins GPX1, GPX4, MSRB1, SELENO H, SELENO K and SELENO S increased under selenium supplementation, while TXNRD1, SEPHS2, SELENO P, N, M and O and did not (Extended Figure 1B). Corresponding mRNA levels did not consistently increase with selenium suggesting this effect is mediated through translation (Extended Figure 1C).

H_2_O_2_ treatment increases Um34 modification in MEFs^30^ and translation of a subset of antioxidant selenoproteins in HEK293T cells^31^. To establish whether oxidative stress has a similar effect in cancer cells, we treated A375 melanoma cells with H_2_O_2_ or a global prooxidant, pyocyanin, which increases intracellular levels of H_2_O_2_ and superoxide^32^, a species of ROS produced by mitochondria complex I and III and released by tumor-infiltrating macrophages and neutrophils^33^. Both H_2_O_2_ and pyocyanin treatment increased Um34 levels. In fact, pyocyanin without selenium increased Um34 to similar levels as selenium supplementation alone (Figure 1D), indicating that oxidative stress drives Um34 biosynthesis independent of selenium status. Oxidative stress increased redox-related selenoproteins (GPX1, GPX4, MSRB1, TXNRD1) in melanoma cell lines (Figure 1E and F) without significant effect on mRNA levels (Figure 1G). Several selenoproteins (SELENOK, SELENOF and SELENOP) were upregulated at the mRNA level suggesting there may be additional transcriptional regulation in response to oxidative stress. Components of the selenocysteine machinery such as SECISBP2 and SEPSECS were not affected by either selenium or oxidative stress.

To measure UGA recoding as a selenocysteine, we established a GFP-Reporter Of Selenocysteine Insertion (gROSI), in which a cysteine71 in the GFP was replaced with a UGA coupled with the 3’UTR GPX4 Selenocysteine Insertion Sequence (SECIS) required for UGA readthrough as a selenocysteine^34^ (Figure 1H). Fluorescence of GFP is dependent upon Sec-insertion at the UGA, which was seen with extracellular selenium (Figure 1I). More importantly, gROSI with either GPX1 or GPX4 3’ UTR SECIS was responsive to oxidative stress induction in the absence of selenium (Figure 1J). The combination of selenium and oxidative stress had an additive effect on Sec-insertion, indicating that Um34 may coordinate selenium and oxidation status to increase selenoprotein translation.

### FTSJ1 is the Um34 selenocysteine-tRNA methyltransferase

To identify the Um34 methyltransferase, we identified proteins interacting with selenocysteine-specific elongation factor EEFSEC in A375 melanoma cells by proteomics. Out of 15 RNA-modifying enzymes interacting with EEFSEC, FTSJ1 was the only wobble 2’O-ribose-modifying tRNA methyltransferase (Figure 2A and Supplementary Table 2)^35, 36^. We validated FTSJ1 interaction with selenocysteine translation machinery by co-immunoprecipitation of FLAG-tagged FTSJ1 with HA-tagged EEFSEC and SECp43 from A375 melanoma cells (Figure 2B, Extended Figure 2A).

**Figure 2:**
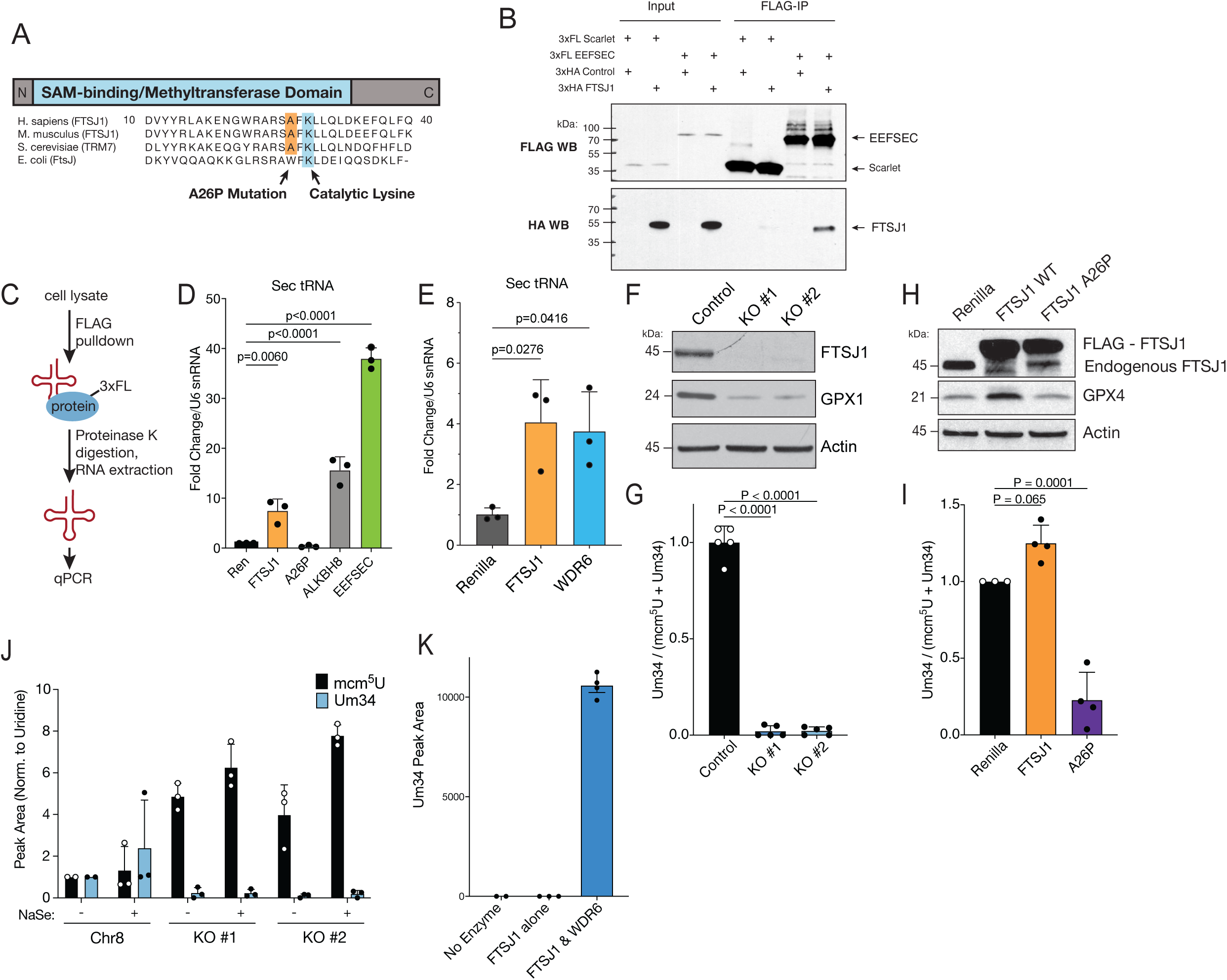
FTSJ1 is the Um34 methyltransferase for tRNA^Sec^. (A) Alignment schematic of FTSJ1 and its homologs (B) Co-immunoprecipitation assay of the interaction between HA-tagged FTSJ1 and FLAG-tagged EEFSEC, compared to FLAG-tagged Scarlet-expressing control vector co-expressed in A375 melanoma cells. Representative from 3 independent experiments shown. (C) Schematic of RNA-immunoprecipitation. (D) Levels of tRNA^Sec^ bound to FLAG-tagged Renilla, wild type and A26P-mutant FTSJ1, ALKBH8 and EEFSEC by qRT-PCR. n = 3. Mean +/- SD of 3 independent replicates. Two-sided unpaired t-test. (E) Levels of tRNA^Sec^ bound to FLAG-tagged Renilla, FTSJ1 and WDR6 by qRT-PCR. n=3 independent replicates. Mean +/- SD of 3 independent replicates. Two-sided unpaired t-test. (F) Western blot analysis of FTSJ1 and selenoprotein GPX1 levels in control and *FTSJ1* knockout A375 melanoma cell lines. Representative from 5 independent experiments shown. (G) Quantification of Um34 modification levels on tRNA^Sec^ from control and *FTSJ1* knockout A375 melanoma cell lines measured by LC-MS/MS. Mean +/- SD. n = 3 independent biological experiment. One-way analysis of variance (ANOVA) followed by Dunnett’s tests for multiple comparisons. (H) Western blot analysis of overexpression of control Renilla luciferase, wild type and mutant A26P FTSJ1 and selenoprotein GPX4 in A375 melanoma cell lines. Representative from 3 independent biological experiments shown. (I) Quantification of Um34 modification levels on tRNA^Sec^ in A375 melanoma cells overexpressing control Renilla luciferase, wildtype and A26P mutant FTSJ1 measured by LC-MS/MS. Mean +/- SD. n = 3-4 independent replicates. Statistical significance assessed by one-way analysis of variance (ANOVA) followed by Dunnett’s tests for multiple comparisons. (J) Quantification of Um34 and mcm^5^U levels on tRNA^Sec^ in Control and *FTSJ1*-KO A375 melanoma cells grown in DMEM/F12 medium with and without selenium (30nM) measured by LC-MS/MS. Mean +/- SD. n=2 independent biological experiments. (K) Quantification of Um34 levels on tRNA^Sec^ in an in vitro methyltransferase assay measured by LC-MS/MS. Mean +/- SD. n= 2 independent biological experiments.

To determine FTSJ1 binding to tRNA^Sec^, we performed RNA-immunoprecipitation from A375 melanoma cells, using either wildtype or catalytically dead A26P mutant of FTSJ1 (found in patients with X-linked intellectual disability) or known tRNA^Sec^-interacting proteins, ALKBH8 and EEFSEC as positive controls (Figure 2C). We saw significant enrichment in the binding of wildtype FTSJ1 to tRNA^Sec^, which was not observed in mutant A26P (Figure 2D). WDR6, the RNA-adapting partner of FTJS1 also showed enrichment for tRNA^Sec^ binding (Figure 2E). FTSJ1 did bind its known substrate tRNA^Phe(GAA)^ but did not show enrichment in binding to other tRNAs such as tRNA^Lys(UUU)^ and tRNA^Glu(CUC)^ (Extended Figure 2B and C).

To functionally validate FTSJ1, we knocked out *FTSJ1* (*FTSJ1*-KO) in A375 (Figure 2F), SK-MEL28 and SK-MEL2 melanoma cell lines. LC/MS-MS analysis of tRNA^Sec^ showed near-complete loss of Um34 on tRNA^Sec^ in the knockout cells compared to control (Figure 2G and Extended Figure 2D and E). Overexpression of wildtype FTJS1 increased Um34 levels, while catalytically inactive mutant A26P had a dominant negative effect, causing a decrease in Um34 levels (Figure 2H and I and Extended Figure 2F). Overexpression of wildtype FTSJ1 in the knockout cell lines rescued the loss of Um34 levels while mutant A26P had no effect (Extended Figure 2G and H). Finally, extracellular selenium had no effect on Um34 levels in the absence of FTSJ1 demonstrating that FTSJ1 activity is the limiting step of Um34 synthesis (Figure 2J). In fact, there was a marked increase in mcm^5^U levels, suggesting a compensatory mechanism by ALKBH8 which generates the mcm^5^U precursor. To definitively show FTSJ1 is the tRNA^Sec^ Um34 methyltransferase, we performed an *in vitro* methyltransferase assay with FTSJ1 and WDR6 on tRNA^Sec^ from *FTSJ1*-KO cells. Um34 levels were only detectable in the presence of both FTSJ1 and WDR6 (Figure 2K and Extended Figure 2I and J). These data directly establish FTSJ1 as the Um34 methyltransferase, demonstrating that FTSJ1 activity is necessary and sufficient for tRNA^Sec^ Um34 modification.

### Um34 loss affects translation of a subset of selenoproteins

To systematically assess changes in protein translation in *FTSJ1*-KO cells, we used ribosomal profiling to quantify ribosome-protected fragments (RPFs) across the transcriptome. The density of RPFs positioned at specific codons indicates the average ribosomal dwelling time and reflects the relative ribosome translation speed across each codon^37, 38^. Given that FTSJ1 is known to modify other tRNAs (Phe tRNA^GAA^ and Trp tRNA^CCA^) in addition to Sec tRNA, we investigated whether loss of FTSJ1 affects translation of individual codons. We compared ribosomal P-site occupancy in *FTSJ1*-KO and Control A375 melanoma cells in the presence of selenium (30nM) and saw that there was no significant difference in codon usage across the global transcriptome especially for codons coding for Phe(GAA) and Trp(CCA) (Extended Figure 3A). This confirms that while FTSJ1 may lead to loss of modifications on other tRNAs, this does not lead to global defects in translation in melanoma cells.

**Figure 3:**
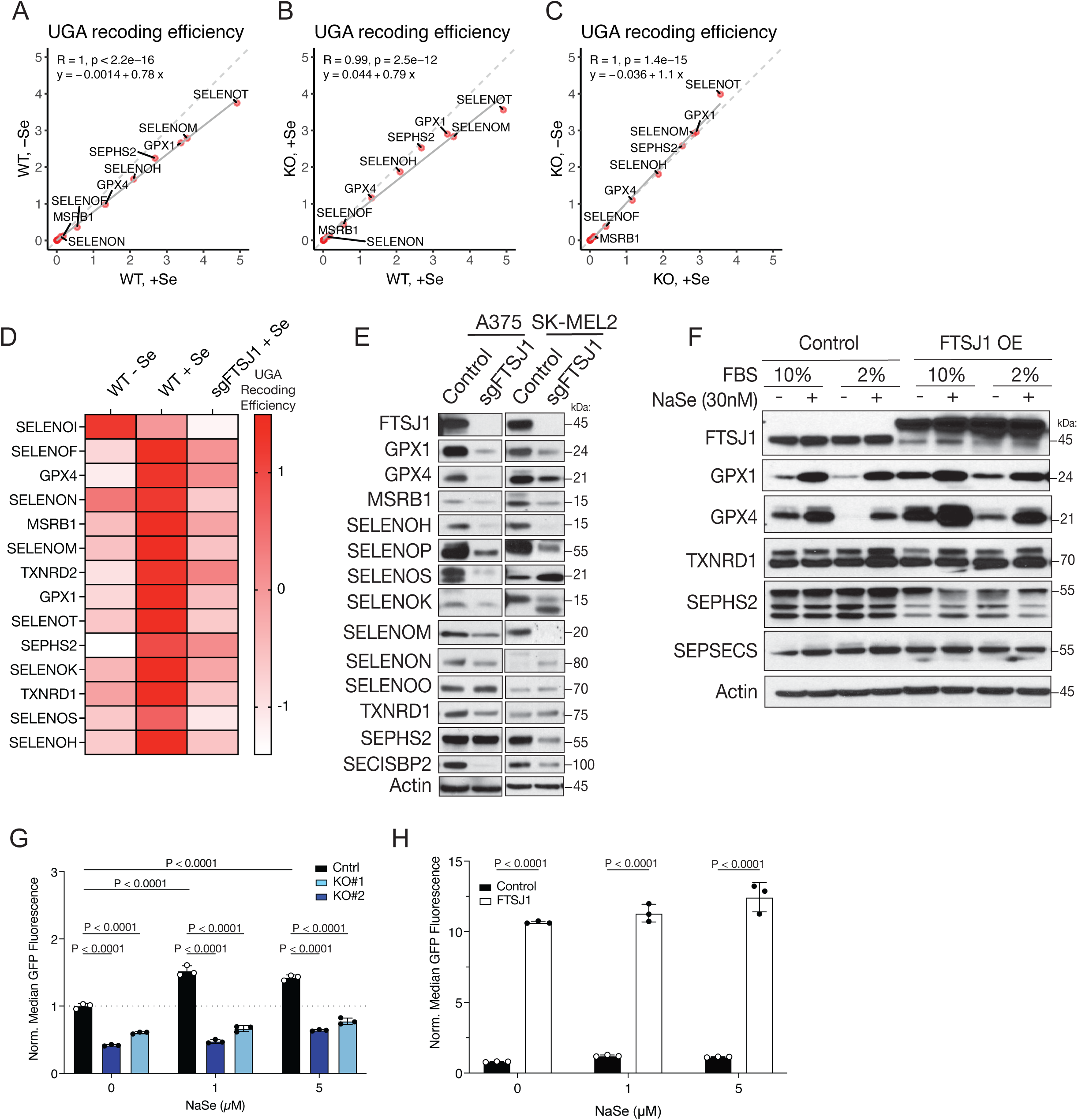
Um34 regulates UGA recoding efficiency in a subset of selenoproteins involved in oxidative stress. (A) UGA Recoding Efficiency (Ratio of RPF 3’ to Sec UGA/RPF 5’ to Sec UGA) of selenoproteins in WT A375 melanoma cells in the absence and presence of selenium (30nM NaSe). n=3 independent replicates per group. Pearson Correlation. (B) UGA Recoding Efficiency of selenoproteins in Control and *FTSJ1*-KO A375 melanoma cells in the presence of selenium (30nM NaSe). n=3 independent replicates per group. Pearson Correlation. (C) UGA Recoding Efficiency of selenoproteins in *FTSJ1*-KO A375 melanoma cells in the absence and presence of selenium (30nM NaSe). n=3 independent replicates per group. Pearson Correlation. (D) Z-score values of UGA recoding efficiency comparing WT A375 melanoma cells without selenium, WT A375 melanoma cells with selenium (30nM NaSe) and *FTSJ1*-KO A375 melanoma cells with selenium (30 nM NaSe). n=3 independent replicates per group. (E) Western blot analysis of selenoprotein levels in Control and *FTSJ1*-KO A375 and SK-MEL-2 melanoma cells grown in normal (10% FBS) media. Representative from 3 independent biological experiments shown. (F) Western blot analysis of selenoproteins in normal (10% FBS), selenium-depleted (2% FBS) and supplemented conditions (10% + 30nM NaSe and 2% + 30nM NaSe) in A375 melanoma cell line overexpressing either control Renilla or FTSJ1. Representative from 2 independent biological experiments shown. (G) Flow cytometry analysis of seleno-GFP reporter (gROSI) expression in control and *FTSJ1*-KO A375 melanoma cells with and without selenium (30nM). Mean +/- SD of 3 independent replicates; representative experiment shown; One-way analysis of variance (ANOVA) followed by Dunnett’s tests for multiple comparisons (two-sided). (H) Flow cytometry analysis of seleno-GFP reporter (gROSI) expression in control Renilla-overexpressing or FTSJ1-overexpressing A375 melanoma cells. Mean +/- SD of 3 independent biological experiments; representative experiment shown; One-way analysis of variance (ANOVA) followed by Dunnett’s tests for multiple comparisons (two-sided).

If tRNA^Sec^ levels are low and/or the ability to decode UGA is impaired (for instance in *FTSJ1*-KO cells), then ribosomes will accumulate upstream of the UGA and translation will terminate with depletion of ribosomes downstream of the UGA. UGA recoding efficiency is defined by the ratio of RPF 3’ to 5’ of the Sec UGA. Comparing UGA recoding efficiency in WT A375 melanoma cells in the presence (30nM NaSe) vs absence of selenium, we saw a significant decrease in recoding efficiency when selenium was depleted (Figure 3A). When UGA recoding efficiency was measured in WT vs *FTSJ1*-KO A375 melanoma cells in the presence of selenium (30nM NaSe), we saw the same decrease in UGA recoding efficiency in *FTJS1*-KO cells (Figure 3B). As a control we compared *FTSJ1*-KO cells in the presence and absence of selenium (Figure 3C), and observed no difference between these two conditions, indicating that FTSJ1-mediated Um34 modification regulates UGA recoding efficiency independent of selenium availability. Z-score was used to compare relative differences in recoding efficiency (Figure 3D). We observed a similar decrease in UGA recoding efficiency in WT A375 melanoma cells without selenium as *FTSJ1*-KO cells, highlighting the role of Um34 modification in UGA readthrough.

We plotted the Cumulative Fraction of RPFs across each selenoprotein transcript to assess ribosomal accumulation. Comparing WT A375 melanoma cells in the presence and absence of selenium (30nM NaSe), we found that a subset of selenoproteins showed an increase in ribosomal accumulation 5’ of the Sec UGA codon when selenium was depleted (Extended Figure 4A). *FTSJ1*-KO cells had a similar ribosomal accumulation 5’ of the Sec UGA codon in this subset of transcripts (Extended Figure 4B). Interestingly, SELENOK was not affected by the absence of selenium in WT A375 melanoma cells but showed significant ribosomal accumulation in *FTSJ1*-KO cells. The opposite was true for SELENON, which showed some accumulation of ribosomes in WT A375 melanoma cells without selenium but was not affected by *FTSJ1*-KO. TXNRD1, TXNRD2, SELENOI and SELENOS were unaffected by either absence of selenium or FTSJ1 loss, however these proteins all contain the Sec UGA codon at the 3’ end of the transcript (Extended Figure 5A and B). Interestingly, when we plotted the non-cumulative RPF counts for each selenoprotein, we saw that loss of FTSJ1 caused accumulation of ribosomes 5’ of the Sec UGA, even on proteins such as TXNRD1 and SEPHS2 which previously appeared to be unaffected by the loss of FTSJ1 (Extended Figure 6). Taken together, these results suggest that FTSJ1-mediated Um34 modification increases the efficiency of UGA readthrough, independent of the specific selenoprotein.

**Figure 4:**
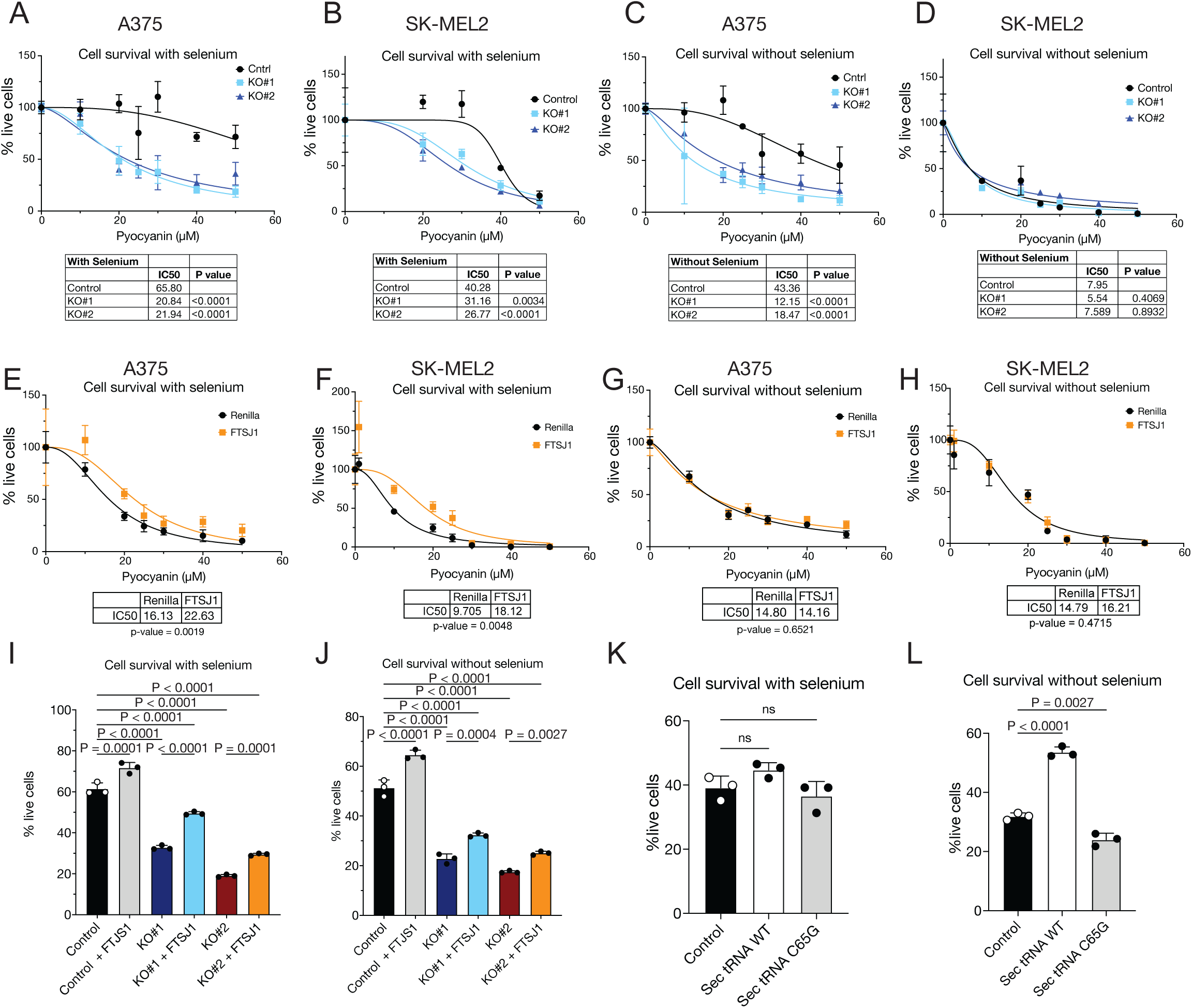
Um34 modification protects cancer cells from oxidative stress. (A-B) Cell survival of Control and *FTSJ1*-KO A375 (A) and SK-MEL-2 (B) melanoma cells under pyocyanin treatment in selenium-supplemented conditions. n = 3 independent culture experiments. Mean +/-SD. Representative of 3 independent experiments. Statistical difference between IC50 values analyzed by Extra-Sum-of-Squares F-Test (two-sided). (C-D) Cell survival of Control and *FTSJ1*-KO A375 (C) and SK-MEL-2 (D) melanoma cells under pyocyanin treatment in selenium-depleted conditions. n = 3 independent culture experiments. Mean +/- SD. Representative of 3 independent experiments. Statistical difference between IC50 values analyzed by Extra-Sum-of-Squares F-Test (two-sided). (E-F) Cell survival of Control and FTSJ1 overexpressing A375 (E) and SK-MEL-2 (F) melanoma cells under pyocyanin treatment in selenium-supplemented conditions. n = 3 independent culture experiments. Representative of 3 independent experiments. Mean +/- SD. Statistical difference between IC50 values analyzed by Extra-Sum-of-Squares F-Test (two-sided). (G-H) Cell survival of Control and FTSJ1 overexpressing A375 (G) and SK-MEL-2 (H) melanoma cells under pyocyanin treatment in selenium-depleted conditions. n = 3 independent culture experiments. Mean +/- SD. Representative of 3 independent experiments. Statistical difference between IC50 values analyzed by Extra-Sum-of-Squares F-Test (two-sided). (I) Cell survival of Control and *FTSJ1*-KO A375 melanoma cells with and without FTSJ1 overexpression treated with 25 uM Pyocyanin in the presence of selenium. n = 3 independent culture experiments. Mean +/- SD. Representative of 3 independent experiments. One-way ANOVA followed by Tukey test for multiple comparisons. (J) Cell survival of Control and *FTSJ1*-KO A375 melanoma cells with and without FTSJ1 overexpression treated with 25 uM of Pyocyanin in the absence of selenium. n = 3 independent culture experiments. Mean +/- SD. Representative of 3 independent experiments. One-way ANOVA followed by Tukey test for multiple comparisons. (K) Cell survival of Control, WT tRNA^Sec^ and C65G-mutant tRNA^Sec^ overexpressing A375 melanoma cells treated with 25uM of Pyocyanin in the presence of selenium. n = 3 independent culture experiments. Mean +/- SD. Representative of 3 independent experiments. One-way ANOVA followed by Dunnett test for multiple comparisons. (L) Cell survival of Control, WT tRNA^Sec^ and C65G-mutant tRNA^Sec^ overexpressing A375 melanoma cells treated with 25uM of Pyocyanin in the absence of selenium. n = 3 independent culture experiments. Mean +/- SD. Representative of 3 independent experiments. One-way ANOVA followed by Dunnett test for multiple comparisons.

**Figure 5:**
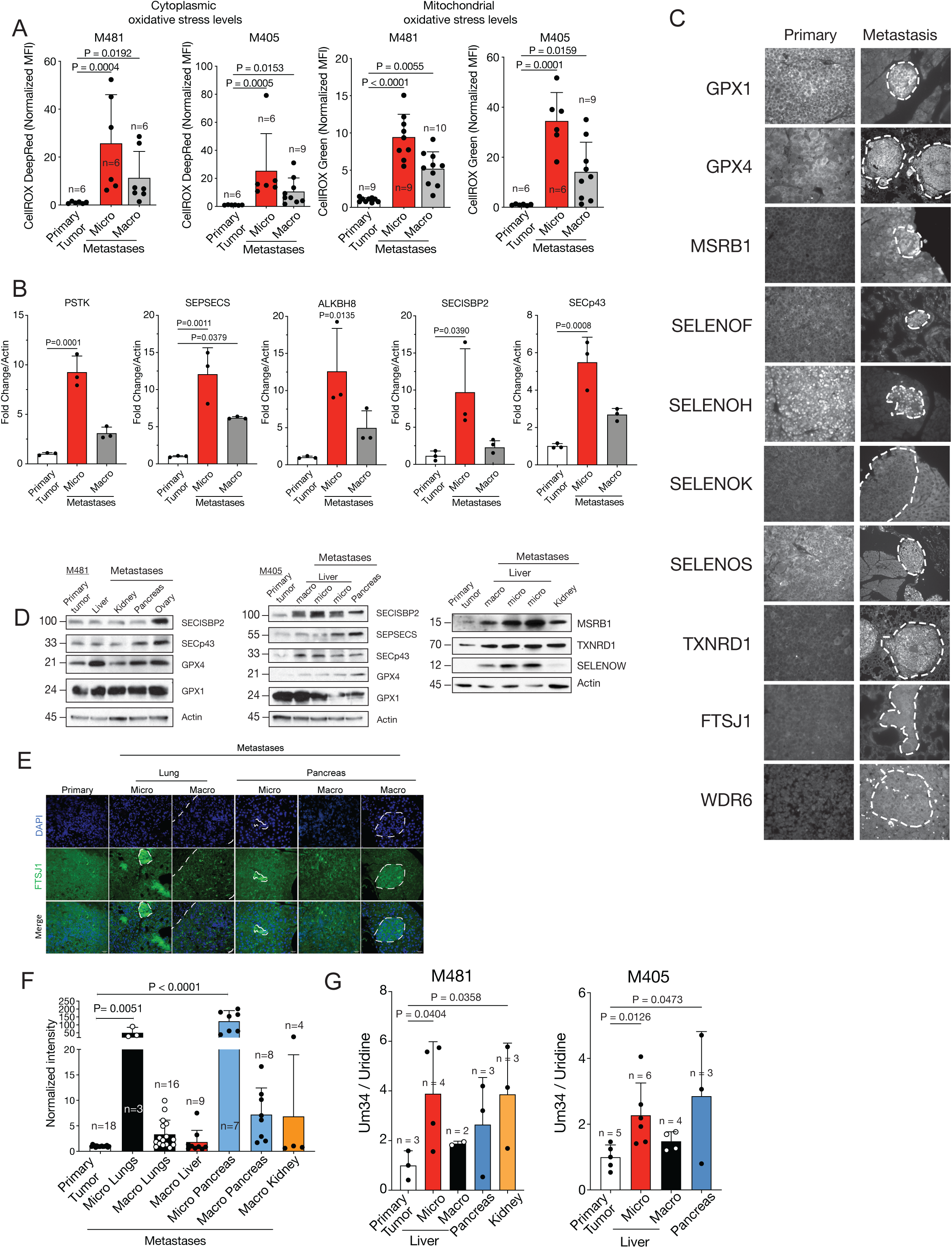
Micrometastases experience high levels of stress and increase selenocysteine biosynthesis. (A) Levels of reactive oxygen species (ROS) in subcutaneous tumors vs micro and macrometastatic nodules from NSG mice transplanted with two different patient melanomas, M481 and M405, measured by flow cytometry with fluorescent dyes. CellROX DeepRed was used to measure cytoplasmic oxidative stress and CellROX Green was used to measure mitochondrial oxidative stress levels. n = number of mice analyzed and indicated in the figure. Mean +/- SD. Statistical significance assessed by one-way analysis of variance (ANOVAs) followed by Dunnett’s tests for multiple comparisons. (B) Transcript levels of selenocysteine biosynthesis machinery in subcutaneous tumors vs micro and macrometastatic nodules from NSG mice transplanted with patient-derived melanoma, M481, measured by qRT-PCR. Mean +/- SD of 3 independent replicates; One-way analysis of variance (ANOVAs) followed by Dunnett’s tests for multiple comparisons. (C) Levels of selenoproteins, FTSJ1 and WDR6 in metastases compared to primary tumor tissues from NSG mice transplanted with patient-derived melanoma, M405, measured by immunofluorescent staining. Representative shown from 3 independent experiments. (D) Levels of selenoproteins and components of selenocysteine biosynthesis machinery in metastases compared to primary tumor from NSG mice transplanted with patient-derived melanomas, M405 and M481, as measured by western blotting. Representative shown from 3 independent experiments. (E) Levels of FTSJ1 in subcutaneous vs micro vs macrometastatic nodules from NSG mice transplanted with Mewo melanoma cell lines, measured by immunofluorescence. (F) Quantification of data shown in (E). n= number of tumors analyzed and indicated in the figure. Mean +/- SD. Two-tailed one sample t-test. (G) Levels of Um34 modification in subcutaneous tumors vs metastatic nodules from NSG mice transplanted with two different patient-derived melanomas, M405 and M481, measured by LC-MS/MS analysis of modified tRNA^Sec^ nucleosides. n= number of mice per melanoma indicated in the figure. Mean +/- SD. One-way analysis of variance (ANOVA) (Kruskal-Wallis test) followed by Dunn’s test for multiple comparisons.

**Figure 6:**
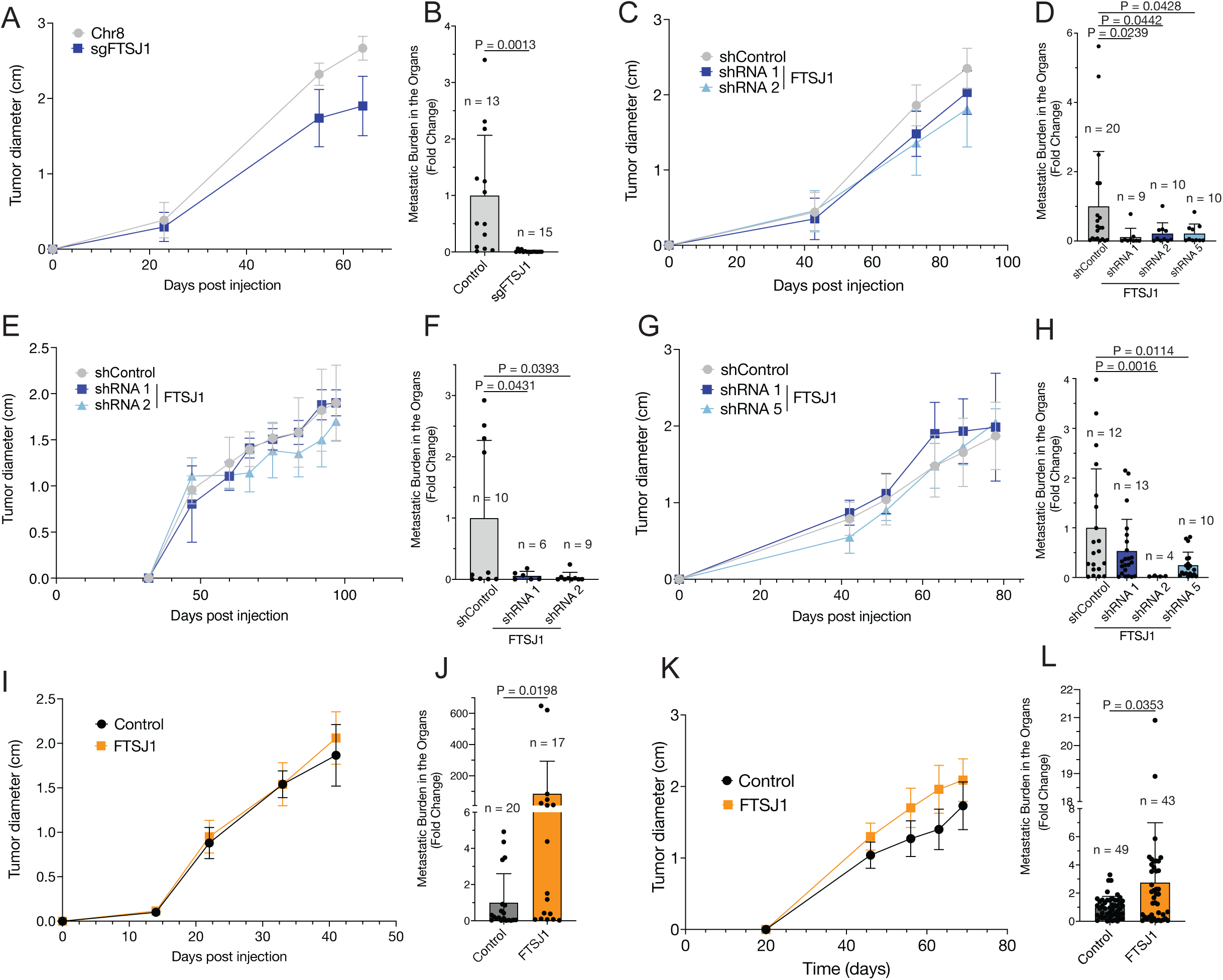
tRNA^Sec^ Um34 modification is necessary and sufficient for metastasis *in vivo*. (A) Subcutaneous tumor growth in mice transplanted with either Control or *FTSJ1*-KO A375 melanoma cells. Data reflect 3 independent experiments but only one representative experiment is shown. Mean +/- SD. For each experiment n= 5 mice. (B) Total metastatic burden in the visceral organs of mice subcutaneously transplanted with either Control or *FTSJ1*-KO A375 melanoma cells. n= number of mice analyzed, indicated in the figure. Mean +/- SD. Two-tailed Mann-Whitney t-test. (C) Subcutaneous tumor growth in mice transplanted with either Control or *FTSJ1* KD MeWo melanoma cells. Data reflect 3 independent experiments but only one representative experiment is shown. For each experiment n= 5 mice. Mean +/- SD. (D) Total metastatic burden in the visceral organs of mice subcutaneously transplanted with either Control of *FTSJ1* KD MeWo melanoma cells. n = number of mice analyzed, indicated in the figure. Mean +/- SD. One-way analysis of variance (ANOVA) followed by unpaired t with Welch’s correction (two-tailed). (E, G) Growth of subcutaneous tumors in mice transplanted with patient-derived melanoma, M405 (E) or M481 (G), expressing either control or *FTSJ1*-targeting shRNAs. For each experiment n= 5 mice. Mean +/- SD. (F, H) Total metastatic burden in visceral organs of mice transplanted with a patient-derived melanoma, M405 (F), or M481 (H) expressing either control or *FTSJ1*-targeting shRNAs. n = number of mice analyzed, indicated in the figure. Mean +/- SD. One-way analysis of variance (ANOVA) followed by unpaired t with Welch’s correction (two-tailed). (I) Subcutaneous tumor growth in mice transplanted with either Control or FTSJ1 overexpressing A375 melanoma cells. Data reflect 3 independent experiments but only on representative experiment is shown. For each experiment n= 5 mice. Mean +/- SD. (J) Total metastatic burden in the visceral organs of mice subcutaneously transplanted with either Control or FTSJ1 overexpressing A375 melanoma cells. n = number of mice analyzed, indicated in the figure. Mean +/- SD. Two-tailed Mann-Whitney t-test. (K) Growth of subcutaneous tumors in mice transplanted from patient-derived melanoma, M481, overexpressing either Renilla luciferase or FTSJ1. Data reflect 3 independent experiments but only on representative experiment is shown. For each experiment n= 5 mice. Mean +/- SD. (L) Total metastatic burden in visceral organs of mice transplanted with patient-derived melanoma, M481, overexpressing either Renilla luciferase or FTSJ1. n = number of mice analyzed, indicated in the figure. Mean +/- SD. Two-tailed Mann-Whitney t-test.

### FTSJ1 regulates Sec insertion independent of selenium

We assessed endogenous selenoprotein levels in two different *FTSJ1*-KO melanoma cell lines, A375 and SK-MEL2. Strikingly, most selenoproteins were decreased in both A375 and SK-MEL2 *FTSJ1*-KO melanoma cell lines (Figure 3E), especially those involved in the antioxidant response (GPX1, GPX4, MSRB1 and SELENOH). SELENOS was only decreased in A375 melanoma cells and SELENOM was only decreased in SK-MEL2 cells, suggesting a difference in selenoprotein regulation depending on genetic background of the cells. Other proteins such as SELENOK, SELENON, SELENOO, TNXRD1 and SEPHS2 were not significantly affected by FTSJ1 loss in either cell line, although SELENOK and SELENON did show a smaller isoform in SK-MEL2 cells which could indicate a truncated protein where the Sec-coding UGA is read as a stop codon. Importantly, these changes were not observed at the mRNA level (Extended Figure 3B), emphasizing that loss of Um34 leads to a translational defect of a subset of selenoproteins. Strikingly, upon loss of FTSJ1 we also saw a decrease in SECISBP2, suggesting FTSJ1 may be involved in stabilizing the SECIS-binding protein complex to promote selenoprotein translation, however this finding warrants further study.

To address whether FTSJ1 is sufficient to drive translation of endogenous selenoproteins when selenium is limiting, we established a system to decrease selenium levels in cells without complete depletion. Due to the variable presence of selenium in serum albumin (150-300nM)^39^, A375 and SK-MEL2 melanoma cells were cultured in either normal (10%) or reduced (2%) FBS levels, with or without selenium supplementation (30nM NaSe). In normal growth conditions (10% FBS, no NaSe), cells had basal levels of selenoproteins GPX1, GPX4 and TXNRD1 (Extended Figure 3C). Growth in 2% FBS, and therefore reduced selenium, led to a decrease in selenoproteins, while selenium supplementation increased selenoprotein levels in both conditions in A375 and HEK-293T cells (Extended Figure 3C). Unlike their proteins, TXNRD1 and GPX4 mRNAs did not change when levels of selenium were varied, suggesting levels of selenium affect translation but not transcription of selenoproteins (Extended Figure 3D). GPX1 mRNA, which is regulated through nonsense mediated decay^40^ showed a modest reduction with reduced FBS but did not increase with selenium supplementation. Compared to control cells, FTSJ1-overexpressing cells had increased antioxidant selenoproteins, GPX1 and GPX4, both in high and low levels of selenium, whereas housekeeping selenoproteins TXNRD1 and SEPHS2 and selenoprotein machinery such as SEPSECS were unaffected by FTSJ1 overexpression (Figure 3F).

To confirm the role of Um34 in selenoprotein translation, we assessed UGA readthrough in *FTSJ1*-knockout cells using the gROSI reporter. Translation of the seleno-GFP was increased in the presence of selenium, but this increase was abolished in *FTSJ1*-knockout cells (Figure 3G). Conversely, overexpression of FTSJ1 caused a significant increase in translation of gROSI independent of selenium levels, indicating that Um34 modification is sufficient for readthrough of UGA as a selenocysteine (Figure 3H). To specifically assess Sec-insertion into selenoproteins, we adapted a method for selenocysteine-specific labeling and enrichment^41, 42^. Selenocysteine residues are labelled with a biotinylated probe at low pH, followed by avidin-immunoprecipitation of the selenocysteine-containing proteins (Sec-IP). As proof of concept, we blotted for known selenocysteine proteins GPX4, SELENOS, SEPHS2 and TXNRD1, in the presence of biotin and selenium, and for the non-selenocysteine containing protein, SECISBP2, as a negative control. (Extended Figure 3E). Next, we performed Sec-IP on Control and *FTSJ1*-KO A375 melanoma cells and found that some selenoproteins, such as GPX1, GPX4, SELENOK, SELENOS were pulled down less in the absence of FSTJ1 (Extended Figure 3F). To confirm that the increase in selenoprotein levels upon FTSJ1 overexpression is due to increased UGA readthrough as a selenocysteine, we performed Sec-IP of control and FTSJ1-overexpressing A375 melanoma cells in the absence and presence of selenium. Even without selenium, FTSJ1-overexpressing cells exhibit as much insertion of Sec into GPX4 and SELENOS as control cells with selenium supplementation (Extended Figure 3G). TXNRD1, traditionally thought to be a housekeeping selenoprotein, did not respond to FTSJ1 overexpression. These data demonstrate that FTSJ1 is both necessary and sufficient for recoding of the UGA as selenocysteine in a subset of selenoproteins and highlight the translational hierarchy that exists among different selenocysteine proteins.

### Um34 protects cancer cells from oxidative stress

Given that Um34 is required for translation of antioxidant selenoproteins and its levels increase in response to oxidative stress, we hypothesize that Um34 and FTSJ1 are required for the oxidative stress response. Surprisingly, there was no increase in oxidative stress levels under normal growth conditions in FTSJ1-deficient melanoma cell lines, and only SK-MEL2 showed increased lipid peroxidation (Extended Figure 7A-C). FTSJ1-deficient cells were more sensitive to prooxidant Pyocyanin both with and without selenium supplementation. However, sensitivity to Pyocyanin greatly increased in both control and *FTSJ1*-KO cells in the absence of selenium, suggesting a crucial role for selenoproteins in the antioxidant response (Figure 4A-D). FTSJ1-deficient cells were also sensitive to ferroptosis, a form of cell death caused by lipid peroxidation, when treated with RSL3 (Extended Figure 7D and E). Overexpression of FTSJ1 protected cells from oxidative stress in the presence of selenium (Figure 4E and F). In the absence of selenium, overexpression of FTSJ1 had no effect, supporting that FTSJ1-driven antioxidant response is dependent on synthesis of selenoproteins (Figure 4G and H). To demonstrate that the sensitivity to oxidative stress was FTSJ1-dependent, we performed rescue experiments and observed that overexpressing *FTSJ1* can partially rescue sensitivity to oxidative stress both in the presence and absence of selenium (Figure 4I and J, Extended Figure 7F and G). FTSJ1 can similarly partially rescue sensitivity of *FTSJ1*-KO cells to ferroptosis induced by RSL3 treatment (Extended Figure 7H and I). Partial rescue is likely due to limiting levels of binding partner WDR6 and insufficient levels of charged tRNA^Sec^. In support of this, overexpression of wild type tRNA^Sec^ increased cell survival under oxidative stress in the absence of selenium (Figure 4K and L and Extended Figure 7J and K), supporting the fact that tRNA^Sec^ may be limiting especially when selenium is depleted. To confirm that this effect is due to the Um34 modification, we overexpressed mutant tRNA^Sec^ C65G - a patient mutation shown to specifically affect translation of stress-related selenoproteins due to loss of Um34 and modifications within the anticodon loop^43,44^. Our data show that tRNA^Sec^ C65G is unable to rescue oxidative stress sensitivity. Consistent with this data, we found that overexpression of wild type Sec tRNA leads to an increase in Um34 levels on tRNA^Sec^ while overexpression of C65G mutant Sec tRNA had no significant effect on Um34 levels (Extended Figure 7L). Finally, to demonstrate oxidative stress sensitivity was not dependent on other FTSJ1-modified tRNAs, we overexpressed either tRNA Phe^GAA^ or Trp^CCA^, and found neither had an effect on oxidative stress resistance with or without selenium (Extended Figure 7M-P, Extended Figure 8A-D). These data confirm that FTSJ1 is both necessary and sufficient for oxidative stress resistance and that this resistance is specific to tRNA^Sec^ containing Um34.

### Micrometastases have higher oxidative stress and Sec biosynthesis

Given the central role of oxidative stress in metastatic colonization, we explored selenoprotein translation in metastasis using a patient-derived xenograft (PDX) model of melanoma metastasis, in which patient melanomas transplanted into NSG mice recapitulate metastatic behavior of the tumor observed in patients^45^. Using two patient tumors with different driver mutations, M481 (BRAF V600E) and M405 (NRAS Q61R), we measured levels of both cytoplasmic and mitochondrial oxidative stress levels in micrometastatic (<0.2mm), macrometastatic (>0.2mm) and primary subcutaneous tumors. Micrometastases had significantly higher levels of oxidative stress in both subcellular compartments compared to macrometastases or primary tumors (Figure 5A). Increase in oxidative stress in micro and macrometastatic lesions was mirrored by a significant increase in the mRNAs for major components of selenocysteine biosynthesis machinery such as PSTK, SEPSECS, ALKBH8, SECISBP2 and SECp43 (Figure 5B). In contrast, we did not see significant transcriptional differences in selenoprotein mRNA levels between micrometastases, macrometastases and primary subcutaneous tumors (Extended Figure 9A, Supplementary Table 3). However, we did observe significant upregulation of certain selenoproteins in metastatic nodules compared to the primary tumors (Figure 5C and D), consistent with the model that selenoprotein translation increases as a protective mechanism against oxidative stress. Overall, our data suggest a translational mechanism for the upregulation of selenoproteins during metastasis. Levels of precursor tRNA^Sec^ were not changing, however levels of mature tRNA^Sec^ were slightly elevated in metastases compared to the primary tumor, similarly suggesting a translational mechanism of selenoprotein upregulation (Extended Figure 9B and C). Levels of FTSJ1 were also elevated in metastases, increasing more in micrometastases compared to macrometastases (Figure 5C and 5E-F). Consistent with FTSJ1 increase, levels of Um34 modification on tRNA^Sec^ were also elevated in metastatic nodules compared to the primary tumor (Figure 5G).

To confirm translational upregulation of selenoproteins in metastasizing melanoma cells, we transplanted A375 melanoma cells expressing the gROSI reporter with no UTR, GPX1 SECIS or GPX4 SECIS. By quantifying GFP signal in metastatic nodules compared to primary tumors, we found that metastases had significantly higher seleno-GFP with either GPX1 SECIS or GPX4 SECIS, demonstrating increased UGA readthrough as a selenocysteine in metastasizing melanoma cells (Extended Figure 9D and E). Overall, these data indicate that metastatic lesions experience higher levels of oxidative stress and have increased selenocysteine biosynthesis, UGA readthrough, and translation of some selenocysteine proteins.

### Um34 promotes metastasis in vivo

To determine if FTSJ1 and therefore tRNA^Sec^ Um34 modification are necessary for metastasis, we transplanted FTJS1 KO A375 melanoma cells co-expressing luciferase subcutaneously into immunocompromised mice and measured both primary tumor growth as well as overall metastatic burden in the organs. Loss of *FTSJ1* lead to significant decrease in metastasis without any impact on the growth of the primary tumor (Figure 6A and B, Extended Figure 10A). MeWo melanoma cells expressing *FTSJ1*-targeting shRNAs exhibited the same phenotype when transplanted *in vivo* with knockdown of *FTSJ1* causing a decrease in metastatic burden without impacting growth of the primary tumor (Figure 6C and D, Extended Figure 10B-D). Depleting FTSJ1 using shRNAs in two different PDX melanoma tumors, M405 and M481, also did not change primary tumor growth but significantly decreased metastatic spread into the organs (Figure 6E-H, Extended Figure 10E and F). A significant change in circulating tumor cell (CTC) frequency was not seen likely due to low levels of CTCs in the PDX model and a potential increased dependence on oxidative stress resistance after metastatic seeding. To address whether FTSJ1 activity was sufficient to promote metastasis, we overexpressed FTSJ1 in A375 melanoma cells and transplanted them *in vivo*. *FTSJ1* overexpression had no effect on primary tumor growth but increased metastatic burden in the organs of the mice (Figure 6I – J, Extended Figure 10G). The same phenotype was observed when *FTSJ1* was overexpressed in a PDX tumor, M481 (Figure 6K-L, Extended Figure 10H). The second PDX tumor, M405, also showed a trend towards an increase in metastasis (Extended Figure 10I - J). To assess whether FTSJ1 had an effect on metastatic colonization, we injected *FTSJ1*-KO cells intravenously and saw a trend towards a decrease in metastasis (Extended Figure 10K). Similarly, when we transplanted FTSJ1 overexpressing cells intravenously, there was a trend towards an increase in metastasis (Extended Figure 10L). We assessed whether levels of lipid peroxidation were specifically affected in either FTSJ1-depleted cells or FTSJ1-overexpressing cells *in vivo*. In either case lipid peroxidation markers did not significantly change in the primary tumor (Extended Figure 10M and N). These data demonstrate that FTSJ1 is both necessary and sufficient for metastatic colonization *in vivo*.

## DISCUSSION

Our findings reveal that Um34, a single modification of the anticodon loop on the selenocysteine tRNA, is required for efficient translation of selenoproteins, promoting oxidative stress resistance and metastasis. We specifically identify oxidative stress as an inducer of Um34 and selenoprotein translation independent of selenium. We characterized FTSJ1 as the Um34 methyltransferase by demonstrating its activity *in vitro*, showing its direct interaction with tRNA^Sec^ as well as selenocysteine translational machinery. Ribosomal profiling data confirm Um34 is required for efficient readthrough of the UGA stop codon on all selenoproteins regardless of selenium status. We hypothesize the apparent hierarchy of selenoprotein translation may be determined by other factors such as transcript and protein stability and the position of the UGA stop codon, however this warrants further study.

The Um34 methyltransferase has been a long standing unknown and previous work on the mechanistic role of Um34 has relied on a mutant tRNA^Sec^ i^6^A37G, lacking the i6A modification – a prerequisite for Um34 formation^40, 46^. No study has directly uncoupled the function of i^6^A from that of Um34. Over the course of this work, FTSJ1 was separately proposed to be the Um34 methyltransferase^36^. Intriguingly, there were no clear selenoprotein translational effects in the Ftsj1-KO adult mouse brain^47^. In contrast, we have established a functional role for Um34-mediated translation of selenoproteins in oxidative stress resistance and specifically showed that Um34 modification is a *bona fide* metastatic promoter in xenograft models of melanoma. Importantly, loss of Um34 specifically impacted metastatic colonization likely by increasing oxidative stress resistance but is dispensable for proliferation of cancer cells in the primary tumor.

FTSJ1 is known to modify several other tRNAs, mainly tRNA^Phe^ and tRNA^Trp^ and its pathogenesis has been implicated in X-linked intellectual disability in patients^48, 49^. Previous studies show that constitutive knockout of FTSJ1 in mice leads to loss of anticodon modifications on 11 tRNAs and aberrant brain function but otherwise normal tissue development in the body. Similarly, patients with mutations in FTSJ1 manifest non-syndromic X-linked intellectual disability but show no dysmorphism. This work emphasized that tRNA^Phe(GAA)^ was most significantly affected in the KO mouse brain which correlated with impaired translation at Phe codons and neurological phenotypes but was unaffected in other adult tissues^36^. Our ribosomal profiling data do not show significant changes in Phe GAA codon readthrough in *FTSJ1*-KO melanoma cells, suggesting FTSJ1 activity may have phenotypic tissue specificity. Given the role of FTSJ1 in brain development, it would be interesting to explore selenoprotein translation at different stages of embryonic neurodevelopment. Further studies are needed to explore other physiological contexts in which tRNA^Sec^ Um34 maybe required, including thyroid hormone regulation, inflammation and neurodegenerative pathologies that are also associated with oxidative stress dysregulation.

Given that redox dysregulation is a hallmark to cancer, it is possible that effects of Um34 loss are more pronounced in cancer cells. Most importantly, metastasis may represent a unique process, highly sensitive to perturbation in oxidative stress, and therefore highly dependent on the antioxidant selenoproteome regulated by Um34 for metastatic colonization. We observed sensitivity to RSL3 *in vitro* upon KO of *FTSJ1*; however, levels of lipid peroxidation were not elevated at baseline. Sensitivity may be due to reduced levels of GPX4, the target of RSL3, however they may be sufficient to maintain lipid peroxidation levels at baseline where cells are cultured under optimal conditions. Additional work is needed to determine if *FTSJ1*-KO leads to increased ferroptosis *in vivo*, where the metastatic cascade generates vastly different stress conditions.

Our work highlights Um34 as a regulatory node that specifically controls the antioxidant response through translation of selenoproteins. It will be particularly important to explore the molecular synergy between how selenium levels and oxidative stress regulate Um34-driven translation. We observe an increase in FTSJ1 protein expression in response to oxidative stress and selenium supplementation as well as in metastatic nodules. ALKBH8, which generates the Um34-precursor, mcm^5^U, is also induced upon oxidation^30^. It is possible that together these Sec tRNA-modifying enzymes coordinate the selenium and oxidation status of the cell to promote translation of selenoproteins. We propose that these enzymes have the potential to be potent therapeutic targets for metastatic disease. Although previous reports have identified other tRNAs as drivers of metastasis^50^, these code for common amino acids that are also required by normal tissues and are difficult to drug. In contrast, tRNA modifying enzymes such as FTSJ1 and ALKBH8 may represent effective drug targets with minimal toxicity to normal tissues.

## METHODS

### Ethics statement

This study complies with all relevant ethical regulations. All experiments were performed according to protocols approved by the animal use committees at Weill Cornell Medicine (Protocol 2017-0033).

### Reagents and Equipment

Commercially available reagents were used without further purification.

### Cell lines and culture conditions

A-375 (CRL-1619), SK-MEL-2 (HTB-68), SK-MEL-28 (HTB-72), MeWo (HTB-65), HeLa (ATCC CCL-2) and HEK-293T (ATCC CRL-3216) cell lines were purchased from ATCC. All cells were maintained in DMEM containing 4.5g/L glucose and L-glutamine without sodium pyruvate (Corning 10-017-CV). All media was supplemented with 10% Fetal Bovine Serum (Thermo Fisher Scientific) and 1% Penicillin Streptomycin (Corning). All cell lines were grown at 37°C with 5% CO2 and tested for mycoplasma.

### Plasmid Construction

miR-E shRNA knockdown constructs were generated by the splashRNA program and ordered from Invitrogen as oligos. Hairpins were ligated (T4 Ligase, NEB) into SiREP lentiviral plasmid behind an SFFV promoter and EGFP fluorescent protein using XhoI and EcoRI sites. For overexpression constructs, the coding sequences of specific proteins were ordered as gene fragments (IDT) with an N-terminus Kozack sequence and 3xFLAG or 3xHA tag. (C-terminus tag: FTSJ1, SECp43, N-terminus tag: SBP2, EEFSEC, WDR6). Constructs were ligated (In-Fusion HD, Takara) into pLenti vector behind an EFS promoter using BamHI and NsiI sites. A cleavable P2A site links the protein expression to a fluorescent protein tag. CRISPR knockout sgRNA sequences were generated using Geneious cloning software and ordered as single stranded oligos from IDT. Vector backbones were kindly provided by Lukas Dow, and all plasmids were sequence verified.

### Lentiviral transduction and generation of stable cell lines

For virus production, 0.9µg of plasmid was combined with 1µg of packaging plasmids (0.4µg pMD2G and 0.6µg psPAX2) and transfected into HEK 293T cells using Polyjet (SignaGen) according to manufacturer’s instructions. Fresh media was added the following day and viral supernatants were collected 72hr after transfection and filtered through a 0.45µM filter. Approximately 1 million cells were infected with viral supernatant and 10µg/mL polybrene (Sigma-Aldrich). Cells then underwent antibiotic selection with puromycin, hygromycin or blasticidin (Inviogen) depending on the antibiotic resistance of each plasmid. For CRISPR knockout, cells stably-expressing Cas9 were infected with the gRNA construct. To generate single clones, cells were diluted and plated into a 10cm dish. Glass rings were used to isolate single cells and grow colonies. All single clones were verified by western blot analysis and Synthego genomic DNA sequencing.

### Selenium supplementation and depletion in cell culture

Sodium selenite (VWR) was used to supplement selenium in cell culture. “Full media” constitutes DMEM media with 10% FBS – which contains trace levels of selenium bound to albumin. 2% FBS in DMEM was used to lower selenium levels in media without depleting cells of FBS. To achieve 0nM Se, DMEM was combined with Ham’s F12 nutrient mix (Thermo Scientific) at a 1:1 ratio with no FBS present. 30nM sodium selenite was then added for selenium supplementation in DMEM/F12 media.

### tRNA biotin enrichment

Small RNA was extracted from cells or tissue using RNAzol RT (Molecular Research Center) according to manufacturer’s instructions. tRNA enrichment was performed as previously described^12^. Briefly, M270 magnetic Streptavidin beads were washed then incubated with a biotin-labelled oligo specific to the 3’ end of the tRNA for 30min at RT with gentle shaking. Beads were then washed to remove excess oligo and resuspended in in 6xSSC buffer (Fisher Scientific). 40-60µg small RNA was also resuspended in 6xSSC buffer. The beads and RNA were incubated separately for 5 min at 75°C and then combined for an additional 5 min incubation. The combined beads and small RNA were then incubated for 40 min at RT with gentle shaking. Beads were then washed three times with 3x SSC, two times with 1x SSC, and several times with 0.1x SSC until the absorbance at 260nm of the wash supernatant was 0. tRNA was eluted in 0.1x SSC by incubating the beads with buffer at 65°C for 3-4 min. tRNA was then precipitated by adding 1µL glycogen (Sigma-Aldrich) and 300mM NaOAc to the tRNA elution followed by the addition of 4x volume 100% ethanol. Samples were then stored at −80°C overnight.

### Analysis of nucleoside modification by LCMS

Ethanol-precipitated tRNA was pelleted by centrifugation for 15min at 10,000xg. After washing with 70% ethanol, the pellet was resuspended in 20µL H_2_O. tRNA samples were enzymatically hydrolyzed to nucleosides as previously described^51^. Briefly, tRNA was then boiled at 100°C for three minutes and then immediately placed in an ice water bath to cool. Two units Nuclease P1 (NEB) were added with 1/10 volume 0.1M ammonium acetate pH 5.3 for two hours at 45°C. 0.002 units of venom phosphodiesterase (Sigma Aldridge) were added with 1/10 volume 1M ammonium bicarbonate and incubated for 2 hours at 37°C. 0.5 units alkaline phosphatase (NEB) was then added and incubated for one hour. Once digestion was complete, the full volume was transferred to a 10kDa centrifugal filter (Pall Nanosep OD010C34) to remove enzymes. Columns were centrifuged for 8 min at 14,000xg at 4°C. Digested nucleosides were then used directly for LC-MS analysis on an Agilent ZORBAX SB-C18, 2.1 × 150mm, 3.5µm column. Chromatography and Mass Spectrometry methods were performed exactly as described in Songe-Moller et al, 2010 on an Agilent HPLC 1290 connected to an Agilent QQQ^12^.

### Analysis of GFP selenocysteine reporter by flow cytometry

The selenocysteine insertion reporter consists of a destabilized-GFP (d2GFP) as described by Corish et al^52^ and a selenocysteine insertion element (SECIS) from GPX1 or GPX4. This is followed by an IRES – iRFP in order to normalize GFP signal to active translation levels in the cell. Both cysteines in GFP were either replaced with UGA, UAA or UAG codons to mimic a UGA-selenoprotein or act as an alternate stop codon control. Cells were transduced with lentiviral constructs containing the reporter and cells were then selected with hygromycin (Invivogen) for one week. GFP and iRFP levels were quantified by flow cytometry on an Attune NxT (Invitrogen). Analysis was completed with FlowJo software and GFP signal was normalized to iRFP signal.

### Real-Time PCR Quantification of tRNA and mRNA

tRNA and mRNA were extracted using a classic Trizol-chloroform RNA extraction. Reverse transcription was performed using iScript cDNA synthesis kit (Rio-Rad) as described by manufacturer. cDNA was then diluted and used for qPCR analysis with SYBR Green PCR master mix (Thermo Fisher). IDT online primer design tool was used to generate qPCR primers and UCSC In-Silico PCR database was then used to verify human-specificity of primers. qPCR analysis was performed on an Applied Biosystems QuantStudio 6 Real-Time PCR System. All targets were normalized to Actin or U6 snRNA in a standard ΔΔCt analysis.

### Immunoprecipitation of protein complexes

Cell lines overexpressing 3xFLAG or 3xHA-tagged proteins were grown in 10cm dishes and protein lysates were made with 1% Triton X-100 lysis buffer with DTT and PMSF added. Cell debris and insoluble fraction was removed by centrifugation. Anti-FLAG M2 (Sigma-Aldrich) or Anti-HA (Thermo Fisher) agarose beads were washed with PBS and added to the cell lysates for an overnight incubation on a rotator at 4°C. The following day, the beads were pelleted by centrifugation at 1000xg at 4°C and washed 3 times with BC100 wash buffer (0.5M Tris HCl pH 7.6, 250mM EDTA, 100mM KCl, 10% glycerol). Protein complexes were eluted with 3x-FLAG peptide (Sigma-Aldrich) or HA peptide (Thermo Fisher) diluted to 1mg/mL in TBS. Protein eluates were then submitted for proteomic analysis or run on 4-20% Tris-Glycine polyacrylamide gels for western blot analysis.

### RNA-immunoprecipitation

Cells were lysed in Triton-X as described above, with the addition of 5µL ribonuclease inhibitor, RNaseOUT (invitrogen) per sample to the lysis buffer. Anti-FLAG M2 agarose beads were added to the lysate for overnight enrichment. The following day, beads were washed as described above and protein/RNA complexes were eluted with 1mg/mL 3xFLAG peptide diluted in TBS. Proteinase K (VWR) was used to digest the protein in the eluate and RNA was extracted. Briefly, PK Buffer (100mM Tris-HCl pH 7.5, 50mM NaCl, 10mM EDTA) was incubated with Proteinase K at 37°C for 20 min to digest any potential RNAses present. FLAG bead supernatant was combined with 100µL Proteinase K solution and incubated for 20 min at 37°C. 100µL 7M Urea buffer (7M Urea, 100mM Tris-HCl pH 7.5, 50mM NaCl, 10mM EDTA) was added to the reaction and incubated for an additional 20 min at 37°C. All 37°C incubations were performed on a thermomixer with gentle mixing. For small RNA extraction, 700µL RNAzol RT was added to the reaction and incubated for an additional 20 min at 37°C. Small RNA was extracted as described by manufacturer’s instructions. Small RNA was precipitated with 20µg glycogen using 1M NaOAC/Ethanol extraction. cDNA was synthesized using iScript cDNA Synthesis Kit (Bio-Rad) and RNA bound to protein complexes was analyzed by qPCR.

### Recombinant protein expression and purification

FTSJ1 (C-6xHis) and WDR6 (N-6xHis) coding sequences were codon-optimized for e.coli bacteria and cloned separately into petDuet-1 (purchased from Addgene) using the NcoI site downstream of the first T7 promoter. Plasmids were transformed into One Shot BL21 (DE3) E. coli (ThermoFisher). Colonies were grown in 6mL LB ampicillin overnight. The following day, cells were expanded to 1 L LB and grown until OD = 0.5-0.6. IPTG was added to 0.1mM and cells were shaken overnight at 18°C and 225 RPM to allow for induction of recombinant protein. The following day, cells were pelleted, washed with PBS and resuspended in lysis buffer (1% Triton X-100, 5mM imidazole in PBS) and sonicated 4x 15 sec at 30% duty cycle output 3 with 1 minute on ice between sonication. Lysates were centrifuged to remove insoluble fraction. NaCl was added to supernatant to raise salt to 500mM. 1mL/sample Ni-NTA resin (Qiagen) was first washed with lysis buffer with 500mM NaCl then beads were added to supernatant and incubated with rotation for 40 min at 4°C. Resin was washed first with 40mL high salt wash buffer (10mM Tris pH 7.8, 1% Triton X-100, 50mM imidazole, 1M NaCl) then 20mL low salt wash buffer (0.5M Tris-HCl pH 7.6, 250mM EDTA, 50mM imidazole, 70mM KCl, 10% glycerol). Recombinant proteins were eluted 4 times with 1mL elution buffer (0.5M Tris-HCl pH 7.6, 250mM EDTA, 250mM imidazole, 70mM KCl, 10% glycerol). Protein was concentrated using Amicon 10K centrifugal filters (Millipore). PMSF was added to all buffers including PBS and DTT was also added to the elution buffer. Purification was validated using Coomassie staining and western blot analysis.

### *in vitro* tRNA methyltransferase assay

*in vitro* assays of FTSJ1/WDR6 2’-O-methylation of Sec tRNA was performed as previously described^35^, however Sec tRNA was extracted from mammalian *FTSJ1*-KO cells to provide a tRNA substrate with mcm^5^U_34_ precursor. Reactions were carried out at 37°C for 2hr in a 100µL reaction mixture containing 50mM Tris-HCl pH 7.5, 200mM NaCl, 10mM MgCl_2_, 100µg/mL BSA, 5mM DTT, 5µM Sec tRNA, 100µM SAM with 1µM FTSJ1 and 0.2µM WDR6. tRNAs were extracted with phenol/chloroform and ethanol precipitated. tRNAs were digested as described above with Nuclease P1, venom phosphodiesterase and alkaline phosphatase then analyzed using HPLC-MS/MS analysis as described above.

### Selenocysteine-specific protein enrichment

Utilizing the differential pKa values of Sec (5.2) and Cys (8.5), we are able to specifically label Sec residues and block Cys residues in both cell lines and tissues^41, 53^.Cell or tissue were lysed in MES lysis buffer pH 5.6 (50mM MES (Sigma Aldrich), 150mM NaCl, 1mM EDTA, 1% Triton X-100) in the presence of protease inhibitors and 0.1mM Iodoacetyl PEG2 biotin (IodoAPB, Fisher Scientific). Protein was quantified and 2mg input was used for protein enrichment by rotating for 1hr at RT to in 0.1mM IodoAPB to selectively label deprotonated selenocysteine residues. Protein was then precipitated with methanol and the pellet was washed a couple times to remove any remaining IodoAPB. Protein pellets were then resuspended in buffered 8M Urea. 5M iodoacetamide (Sigma-Aldrich) was added for 1hr at RT to selectively block deprotonated cysteines. Proteins in urea were added directly to streptavidin agarose beads (Invitrogen) in PBS, 0.1% SDS and rotated overnight at 4°C. The following day, beads were washed three times with 2M Urea in PBS, 0.1% SDS and then three additional times in 2M Urea in PBS without SDS. For western blot analysis, beads were boiled in 1x protein loading buffer at 95°C for 5 minutes.

### Ribosomal Sequencing

The Ribo-seq libraries were generated as previously described with following modifications (McGlincy et al.)^38^. Cells were snap-frozen and allowed to thaw on ice, then lysed in lysis buffer [50 mM Tris-HCl (pH 7.4), 150 mM NaCl, 5 mM MgCl_2_, 1 mM DTT, 1% Triton X-100, 100 μg/mL cycloheximide, 100 μg/mL tigecycline, 15 U/mL DNaseI (ThermoFisher, 89836)]. RNA was quantified using Qubit Broad range RNA kit (ThermoFisher, Q10211). Ribosome footprint was generated using 15 U RNase I (Biosearch Technologies, N6901K) to 300 ug RNA in 200 μL reaction volume at room temperature for 45 min. The ribosome was collected by ultracentrifugation and ribosome footprint was purified using TRIzol (ThermoFisher, 15596018). RNA was then dephosphorylated and subjected to size selection from 17-40 nt range from 15% polyacrylamide TBE-Urea gel. Ribosome footprint was extracted from the gel in the extraction buffer [50 mM Tris-HCl (pH 7.0), 300 mM NaCl, 1 mM EDTA, 0.25 % SDS] and subjected to rRNA depletion using Human/Mouse/Rat riboPOOL kit (Galen Molecular, dp-K024-000050) by following the manufacture’s instruction. rRNA-depleted ribosome footprint was precipitated in 80% ethanol at −20 °C overnight. rRNA-depleted ribosome footprint was then ligated to a pre-adenylated linker (rApp-WWAGATCGGAAGAGCACACGTC-3ddC). The footprint was reverse transcribed by SuperScript III reverse transcriptase (ThermoFisher, 18080044) with a 5’ phosphorylated primer that contains eight-nucleotide unique molecular identifiers and two spacer residues (Phos-NNNNNNNNAGATCGGAAGAGCGTCGTGTA-iSp18-AA-iSp18-TAGACGTGTGCTC). cDNA was run on 10% non-denaturing gel to remove the non-ligated linker and the reverse transcription primer. cDNA was the circularized using CircLigase I in the presence of 1 mM ATP. The library was PCR amplified using KAPA HiFi HotStart ReadyMix (FisherScientific, 50-196-5217) and NEBNext® Multiplex Oligos for Illumina® Dual Index Primers Set 1 (NEB, E7600S). The library was purified twice using 1x AMPure XP Bead (BeckmanCoulter, A63881). The library was sequenced for single-end 100 cycles on Illumina NextSeq 2000 or paired-end 100 cycles on Illumina NovaSeq 6000.

### Ribo-Seq Analysis

Ribo-Seq reads were trimmed using Trim Galore v0.6.10 (https://github.com/FelixKrueger/TrimGalore/tree/master) to remove nucleotides with low quality and adaptor contamination. Duplicated reads created by PCR amplification were removed using the UMI reads and running fastq2collapse.pl and stripBarcode.pl scripts from CTK tool kit (PMID: 27797762). Then, non-coding RNA were removed using a custom reference genome composed by miRNA, rRNA, tRNA and lncRNA sequences. miRNA sequences were downloaded from RNAcentral database (https://rnacentral.org), tRNA sequences were obtained from GtRNAdb (http://gtrnadb.ucsc.edu/index.html) using High confidence GRCH38/hg38 tRNA sequences, lncRNA were downloaded from LNCipedia (PMCID: PMC3531107) and rRNA reference sequence was built using U13369.1, NR_145819.1, NR_146144.1, NR_146151.1, NR_146117.1, X12811.1, NR_003287.4, NR_023379.1, NR_003285.3 and NR_003286.4. Mapping was performed using STAR v2.7.9a (PMCID: PMC3530905) with default parameters and reads that mapped to the custom reference were removed. FastQC v0.11.8 was used for quality control of the remaining reads. STAR v2.7.9a was used for a second mapping step to the full human reference genome, with – alignEndsType EndToEnd and –quantMode TranscriptomeSAM, using GRCh38 primary assembly genome and MANE v1.2 annotation file to obtain transcriptome and genome mapping coordinates.

Aligned bam files originated from mapping to the transcriptome were used as input for riboWaltz package (https://doi.org/10.1371/journal.pcbi.1006169). Nucleotide base in-frame p-sites coverages were calculated to determinate stalling based on downstream cumulative distribution function (CDF), to identify UGA recoding efficiency by 5’ and 3’ footprints, and to calculate p-site codon usage. For in-frame p-site coverage quantification, reads longer than 25 pb and shorter than 45 pb were kept and p-offsites were calculated for each read length using psite function in riboWaltz with default arguments. Once, p-site position of each read was identified, only in-frame p-sites were kept for further analysis. Mean values and cumulative fractions across the replicates in the different conditions were calculated and plotted using the in-frame p-site coverage in selenoproteins.

In a similar way, UGA recoding efficiency was determined based on the obtained CDS p-site coverage. Ratio of 5’ and 3’ coverages were calculated on fixed -X and +X bases relative to the Sec codon. Since we were using only CDS p-site values and for some of the selenoproteins the distance between Sec and stop codon was short, these fixed bins were different between selenoproteins (refer to the GitHub code to identify the number of bases used for each selenoprotein). Codon usage of p-site was calculated using codon_usage_psite function from riboWaltz. Script used in this analysis are available at https://github.com/abcwcm/piskounova_ribo.

### Tissue RNA-Sequencing

RNA-seq library from primary and metastatic tumors from PDX model of melanoma metastasis was prepared by snap freezing tissue in Trizol. RNA was extracted according to manufacturer’s instructions using TRIzo (ThermoFisher, 15596018) and quantified on NanoDrop. RNA library was generated using TruSeq Stranded Total RNA Library Preparation (rRNA depletion and Stranded RNA-Seq). The library was sequenced for single-end 100 cycles on Illumina NextSeq 2000.

### Tissue RNA-Seq Analysis

Mouse reads from Xenograft data were removed using *bbsplit.sh* from BBMap v38.90 (BBMap - Bushnell B. - sourceforge.net/projects/bbmap/) with ambiguous2==”toss” and using gencode GRCh38 human and GRCm39 mouse references. Using same tools and parameters as for Ribo-Seq analysis, low quality reads and adaptors were trimmed, and non-coding RNA reads were removed. In the same way, the quality control step was performed using FastQC v0.11.8 and a second mapping to the full human reference genome was performed using STAR v2.7.9a and same parameters as in Ribo-Seq. Using bam files of the mapped reads to the whole genome gene counts quantification of CDS regions was calculated using featureCounts v 2.0.1 (PMID: 24227677). Using the obtained counts PCA analysis was performed to analyze the similarities among the samples. This analysis identified significant patient donor effect (M405, M481) and therefore, normalization and further differential expression analysis based on DESeq2 (PMID: 25516281) approach were performed separately for M405 and M481. Wald test was applied to identify changes in Selenoproteins between primary, macro and micro tumor types and adjusted p-values were based on Benjamini-Hochberg procedure.

### Analysis of cell survival under prooxidant treatment

Cells were plated at 10,000 cells/well in a white TC-treated 96 well plate (Corning). The following day, DMEM/F12 media with 0nM or 30nM selenium was added. After 24 hours of selenium depletion or supplementation, cells were then treated with prooxidant pyocyanin (Cayman), H_2_O_2_ (Sigma-Aldrich) or translation inhibitor tunicamycin (Tocris). After 24 or 48 hours of treatment, CellTiter-Glo 2.0 (Promega) was used according to manufacturers instructions to quantify cell viability.

### Lentiviral transduction of human melanoma cells

Patient-derived tumors M405 and M481 were derived as described in^45, 54, 55^. A lentiviral construct with luciferase and dsRed (luc P2A dsRed) was used to label patient-derived melanoma cells as described in^29^. Virus was produced as described above. For lentiviral transduction, 500,000 freshly dissociated melanoma cells were infected with viral supernatant and supplemented with 10 µg/mL polybrene (Sigma). The following day, the media was replaced and 48hrs post-infection cells were either injected subcutaneously into mice as bulk tumors or FACS sorted for positive infection.

### Cell labeling and sorting

All melanoma cells in this study stably express dsRed and luciferase so that melanoma cells could be distinguished by flow cytometry or bioluminescent imaging. When preparing cells for sorting by flow cytometry, cells were stained with antibodies against mouse CD45 (30-F11-VioletFluor, Tonbo), mouse CD31 (390-VioletFluor, eBiosciences), Ter119 (Ter-119-VioletFluor, Tonbo) and human HLA-A, -B, -C (BD Biosciences) to select live human melanoma cells and exclude mouse endothelial and hematopoietic cells. Antibody labelling was performed for 20 min on ice, followed by washing and centrifugation. Before sorting, cells were resuspended in staining medium (L15 medium containing bovine serum albumin (1 mg/mL), 1% penicillin/streptomycin, and 10mM HEPES, pH 7.4) containing 4’6-diamidino-2-phenylindole (DAPI; 5µg/mL; Sigma) to eliminate dead cells from sorting. Live human melanoma cells were isolated by flow cytometry by sorting cells that were positive for dsRed and HLA and negative for mouse CD45, CD31, Ter-119 and DAPI.

### Transplantation of melanoma cells

After sorting, cells were counted and resuspended in staining medium with 25% high-protein Matrigel (product 354248; BD Biosciences). Subcutaneous injections were performed into the right flank of NOD.CB17-*Prkdc^scid^ Il2rg^tm1Wjl^*/SzJ (NSG) mice (Jackson Laboratory) in a final volume of 50 μl. Both male and female mice, 5-8 weeks old were used. Sex was not considered in the study design. Mice were randomized between control and experimental groups, where mice with comparable age and body weight were assigned into control and experimental groups. For a standard metastasis assay, 100 cells were injected subcutaneously into the right flank of the mice. Tumor formation was evaluated by regular palpitation of the injection site and tumors were measured weekly until any tumor in the mouse cohort reached 2.5cm in its largest diameter. Mice were monitored daily for signs of distress according to a standard body condition score or within 24hr of their tumors reaching 2.5 cm in diameter – whichever came first. These experiments were performed according to protocols approved by the Institutional Animal Care and Use Committee at Weill Cornell Medicine (protocol 2017-0033). If tumor burden was found to exceed 2.5cm, mice were euthanized immediately as was allowed by the IACUC protocol 2017-0033. For intravenous transplantation, 1000 A375 melanoma cells were injected into the tail vein of NSG mice in 100ul of PBS. Mice were imaged weekly by bioluminescence until signal was saturated.

### Animal studies

For bioluminescent imaging, mice were injected intraperitoneally with 100µL of DPBS containing D-luciferin monopotassium salt (40µg/mL, Goldbio) 5 min before imaging, followed by general anesthesia with isoflurane 2 min before imaging. IVIS Imaging System 200 Series (Caliper Life Sciences) with Living Image Software was used with the exposure time set to 10 seconds. After imaging the whole body, mice were euthanized and individual organs were dissected and imaged. The bioluminescent signal was quantified with ‘region of interest’ measurement tools in Living Image (Perkin Elmer) software. After imaging, tumors and organs were fixed in paraformaldehyde for histopathology. Studies received ethical approval by animal use committees at Weill Cornell Medicine (2017-0033).

### Circulating tumor cell analysis

During imaging described above, once mice were euthanized for organ dissection, blood was also collected by cardiac puncture with a syringe pretreated with citrate-dextrose solution (Sigma). Red blood cells were precipitated by Ficoll sedimentation according to manufacturer’s instructions (Ficoll Paque Plus, GE Healthcare). Remaining cells were washed with HBSS (Invitrogen) before antibody staining as described above and flow cytometric analysis.

### Immunofluorescence staining of tissue sections

Tissues were fixed in 4% paraformaldehyde for 12 h at 4 uC, embedded in paraffin, and then sectioned. Sections (10 mm) were deparaffinized and then subjected to antigen retrieval for 10 minutes at 121 uC at 15 PSI in 2.94 g/L sodium citrate (pH 6; Sigma Aldrich, 61332-04-3). Sections were permeabilized in PBS with 0.4% Triton X-100 (PBT) and 1% bovine serum, for 10 min, and blocked in 5% bovine serum in PBT for 45 min at room temperature. Sections were then stained with primary antibodies overnight at 4uC: FTSJ1 (A7967, ABclonal; 1:250), GPX1 (A11166, ABclonal; 1:250), GPX4 (52455S, Cell Signaling; 1:250), MSRB1 (A6737, ABclonal; 1:250), Seleno F (SEP15) (A7967, Abcam; 1:250), SELENOH (HPA048362, Atlas Antibodies; 1:250), SELENOK (HPA008196, Atlas Antibodies; 1:250), SELENO S (VIMP-1) (15160S, Cell Signaling Technology; 1:250), TXNRD1 (15140S, Cell Signaling Technology; 1:250), GFP (13970, abcam, 1:1000) in PBS with 0.1% Triton X-100 and 1% bovine serum. The next day, sections were washed in PBS with 0.1% Triton X-100 and 1% bovine serum and stained with secondary, either goat anti-rabbit antibody (A-21244, Invitrogen; 1:500) or goat anti-chicken (ab150169, abcam; 1:1000), for 1 hour in the dark at room temperature. Sections were washed with PBS with 0.1% Triton X-100 and 1% bovine serum with DAPI (1:1500) for four minutes in the dark at room temperature and mounted for imaging. Fluorescent signal was analyzed and quantified using QuPath v0.4.3 software.

### Statistics and Reproducibility

No statistical methods were used to predetermine sample size but we used adequate numbers of samples that would provide statistically significant results based on our previous experience with these tumor models. The data in all figure panels reflect several independent experiments as indicated in figure legends. Variation is always indicated using standard deviation. For analysis of statistical significance, when only two conditions were compared a two-sided unpaired t-test was used, unless otherwise indicated. For comparison of multiple conditions, one-way ANOVA was used for multiple comparisons, unless otherwise indicated. P value of <0.05 was considered statistically significant. Data was assumed to be normally distributed, but this was not formally tested. Individual data points are shown where appropriate. All *in vivo* experiments were randomized to each experimental cohort. The number of mice used is indicated in figure panels. Any mice that died before the end of the experiment due to opportunistic infection, the data from these mice were excluded. No blinding was used in any experiment. Otherwise no data was excluded from the analyses. Further information on research design is available in the Nature Research Reporting Summary linked to this article.

## Data Availability

Ribosomal Sequencing and RNA-sequencing data that support the findings of this study have been deposited in the Gene Expression Omnibus (GEO) under accession codes GSE270971 and GSE270976. Mass spectrometry data for Proteomics as well as nucleoside modification analysis have been provided as Supplementary Data files. Source data for have been provided as Source Data files. All other data supporting the findings of this study are available from the corresponding author on reasonable request by email. Requests will be processed within 30 days.

## Code availability

All custom scripts used in this study are deposited at https://github.com/abcwcm/piskounova_ribo

## ACKNOWLEDGEMENTS

This work was supported by American Cancer Society (EP), Elsa U. Pardee Foundation Grant (EP) and Feldstein Medical Research Foundation Award (EP). LN is supported by Ruth L. Kirschstein Predoctoral Individual National Research Service Award.

## AUTHOR CONTRIBUTIONS

LN designed and performed experiments, analyzed data, and wrote the paper. KC, ID, MZ, GC, RH and KA performed experiments and analyzed data. LED supervised CRISPR/Cas9 experiments and provided reagents. SM and SJ designed, performed and supervised Ribo-Seq experiments. MA, PZ and DB designed, performed and supervised Ribo-Seq and RNA-seq data analysis. EP designed, performed, and supervised experiments, analyzed data, supervised data analysis, and wrote the paper.

---

**Extended Figure 1.**
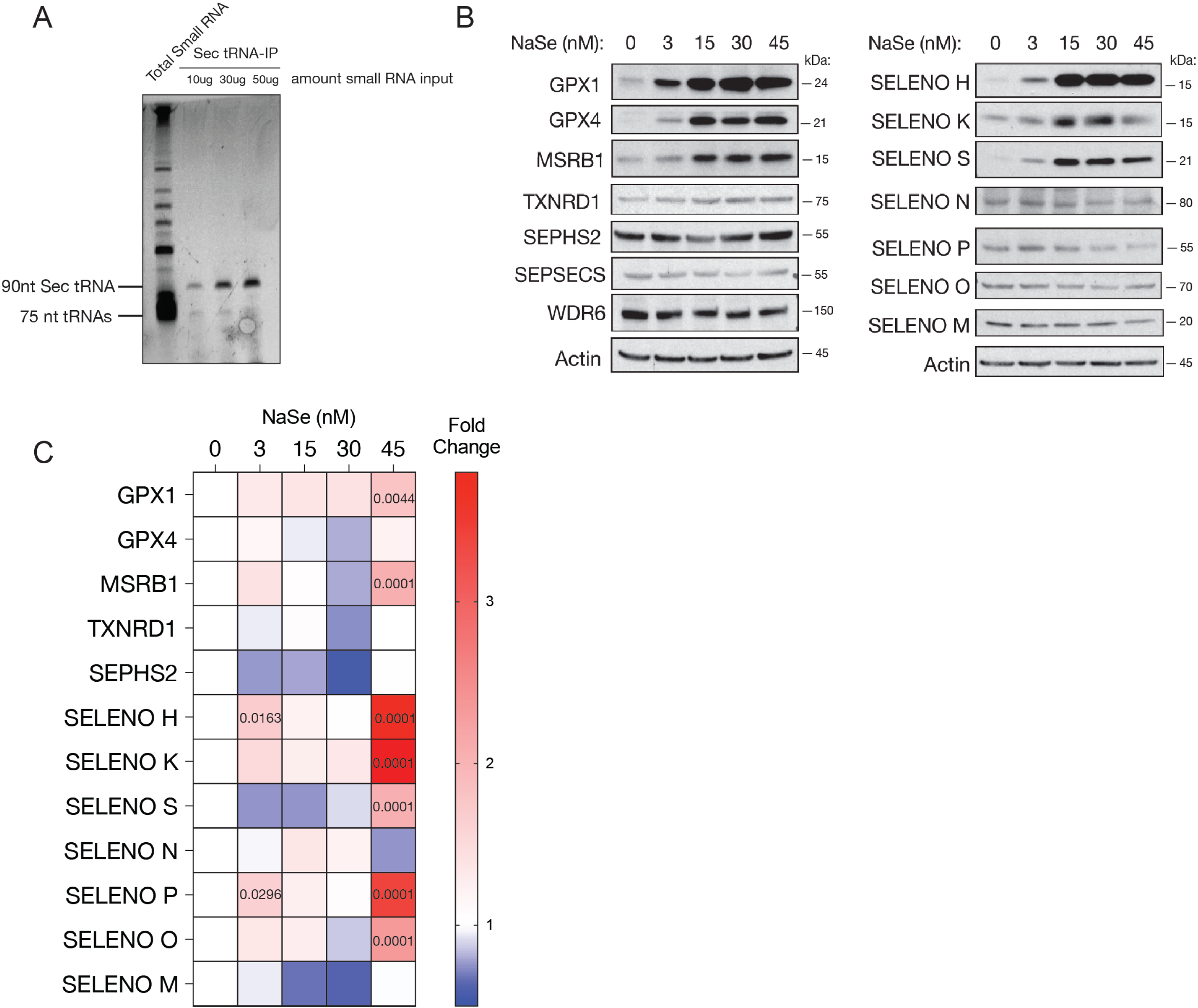

**Extended Figure 2.**
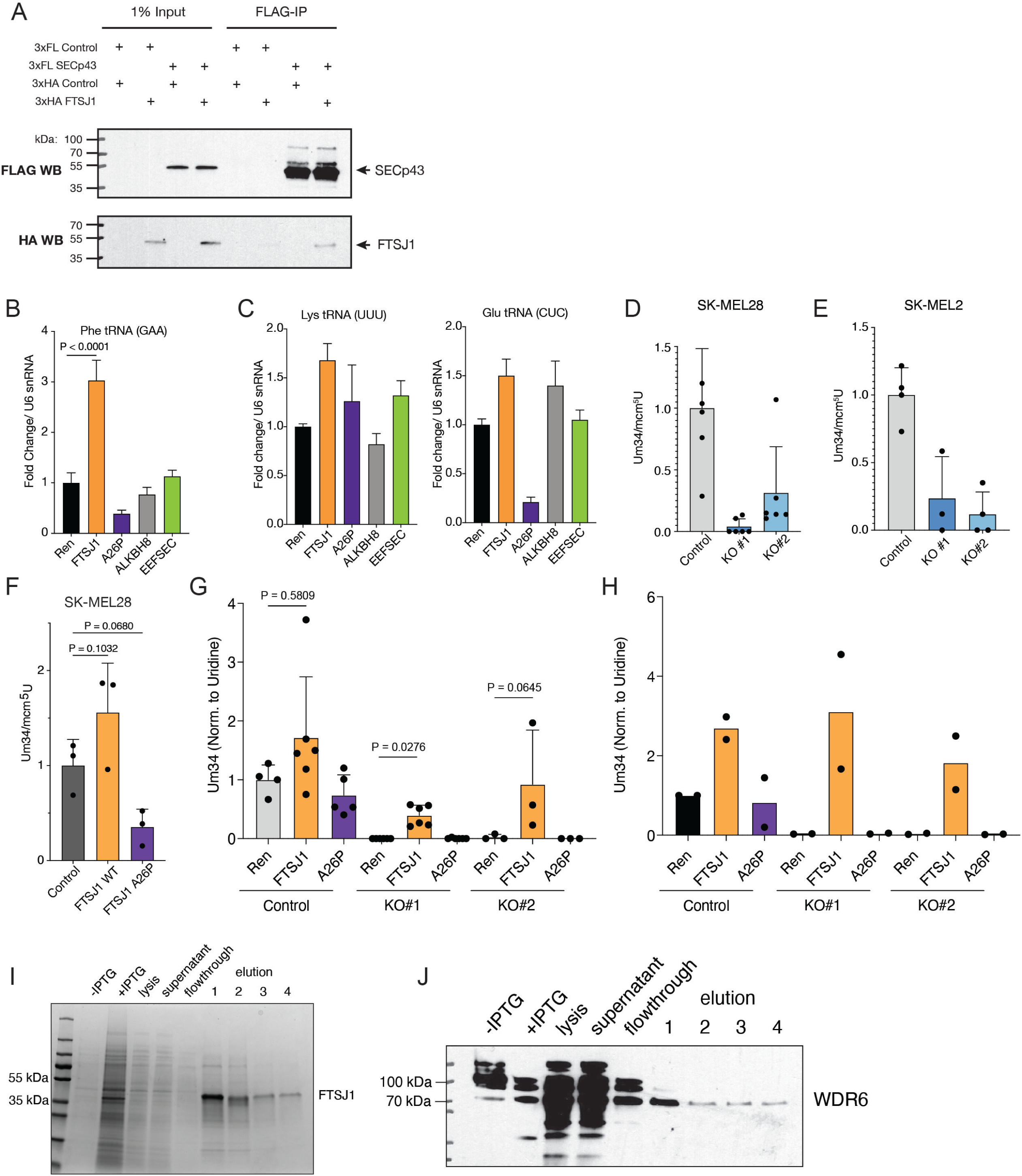

**Extended Figure 3.**
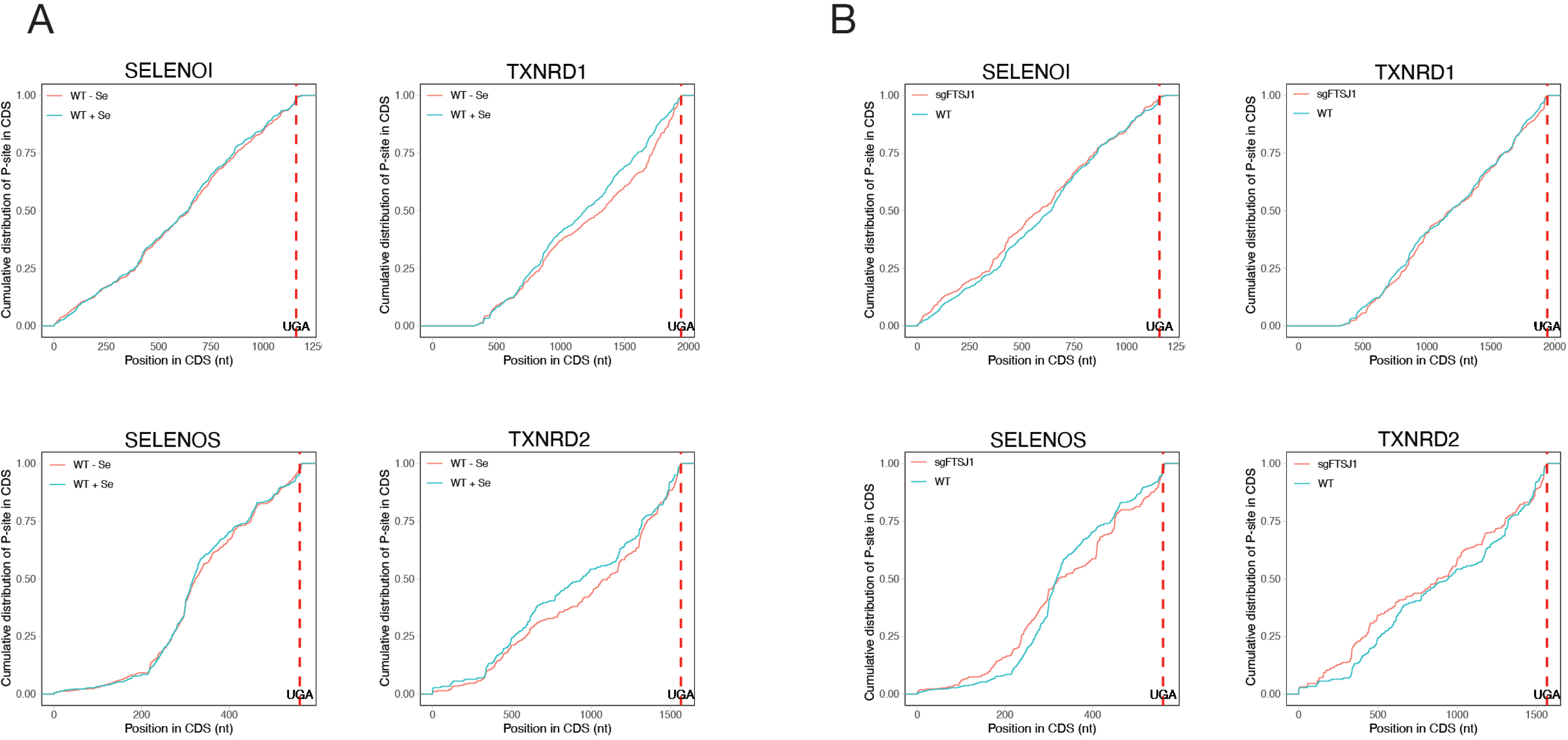

**Extended Figure 4.**
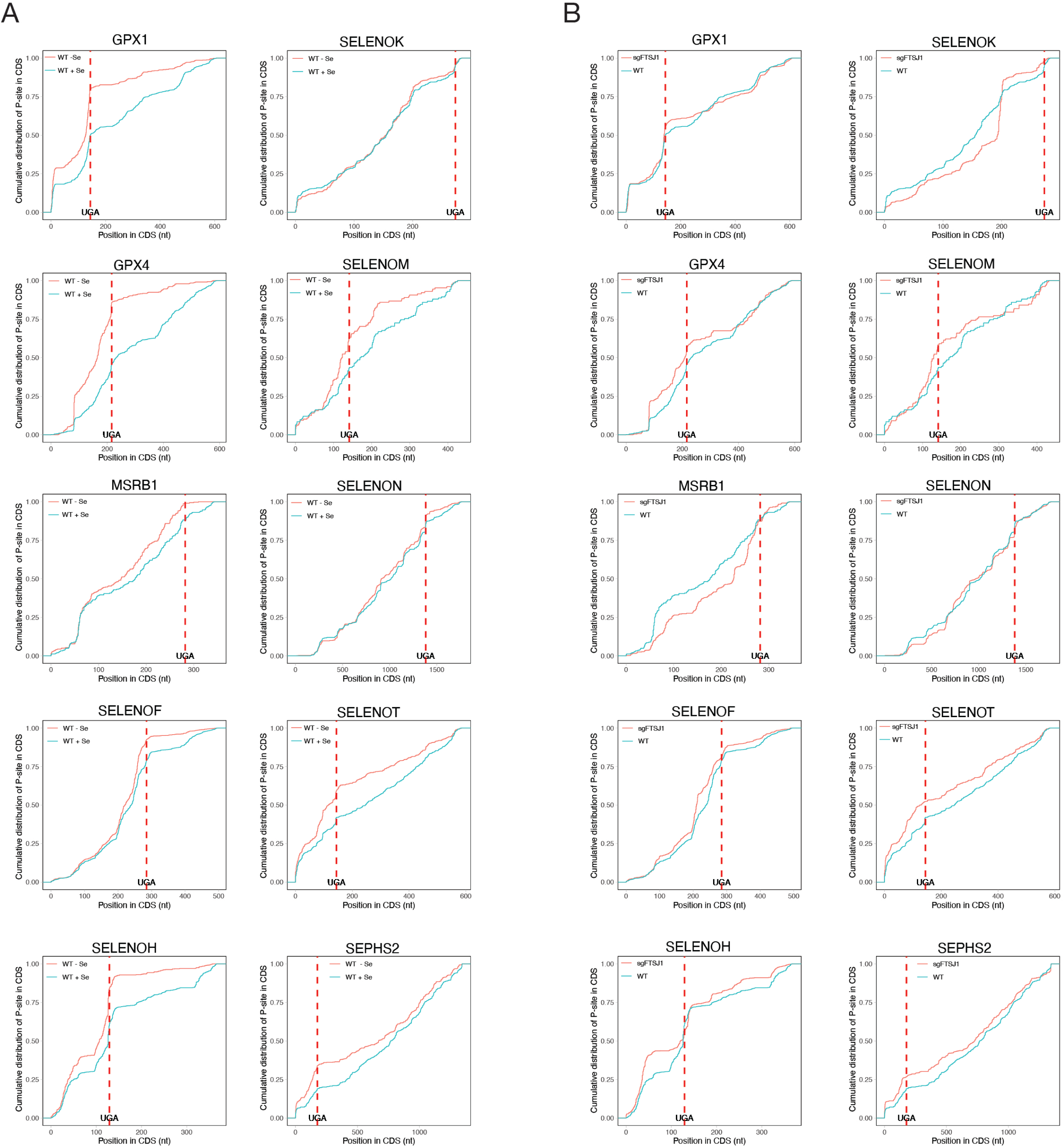

**Extended Figure 5.**
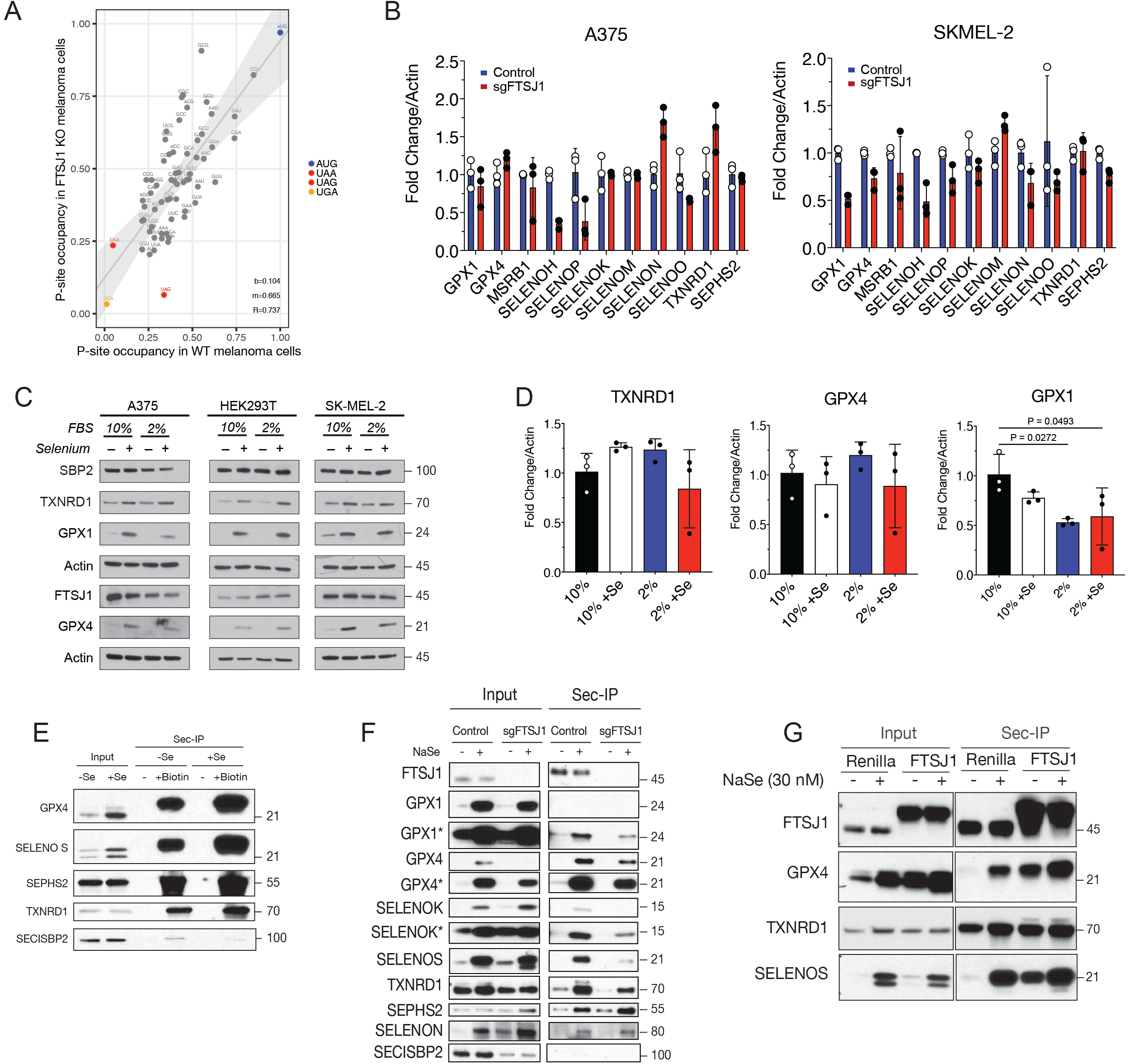

**Extended Figure 6.**
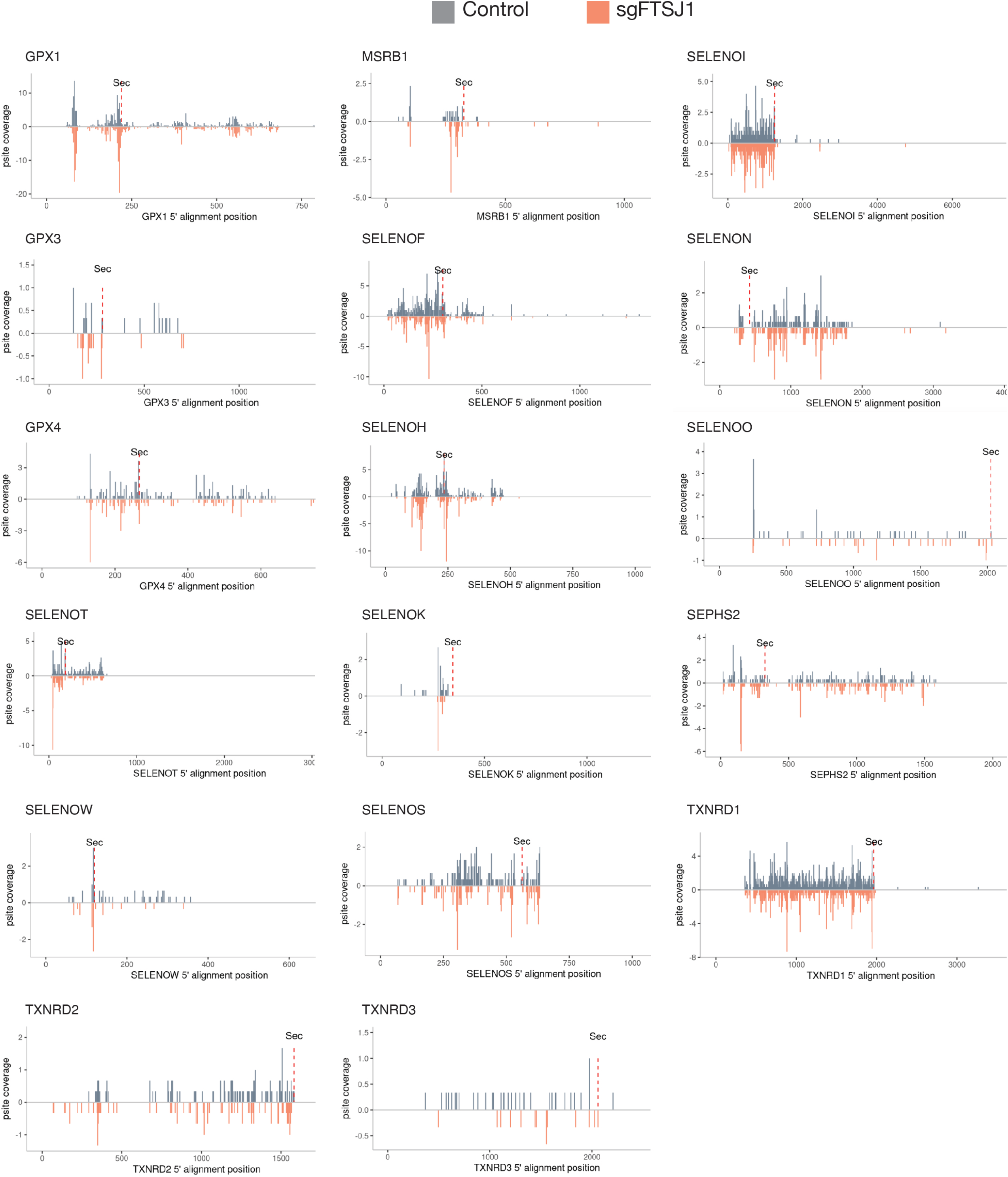

**Extended Figure 7.**
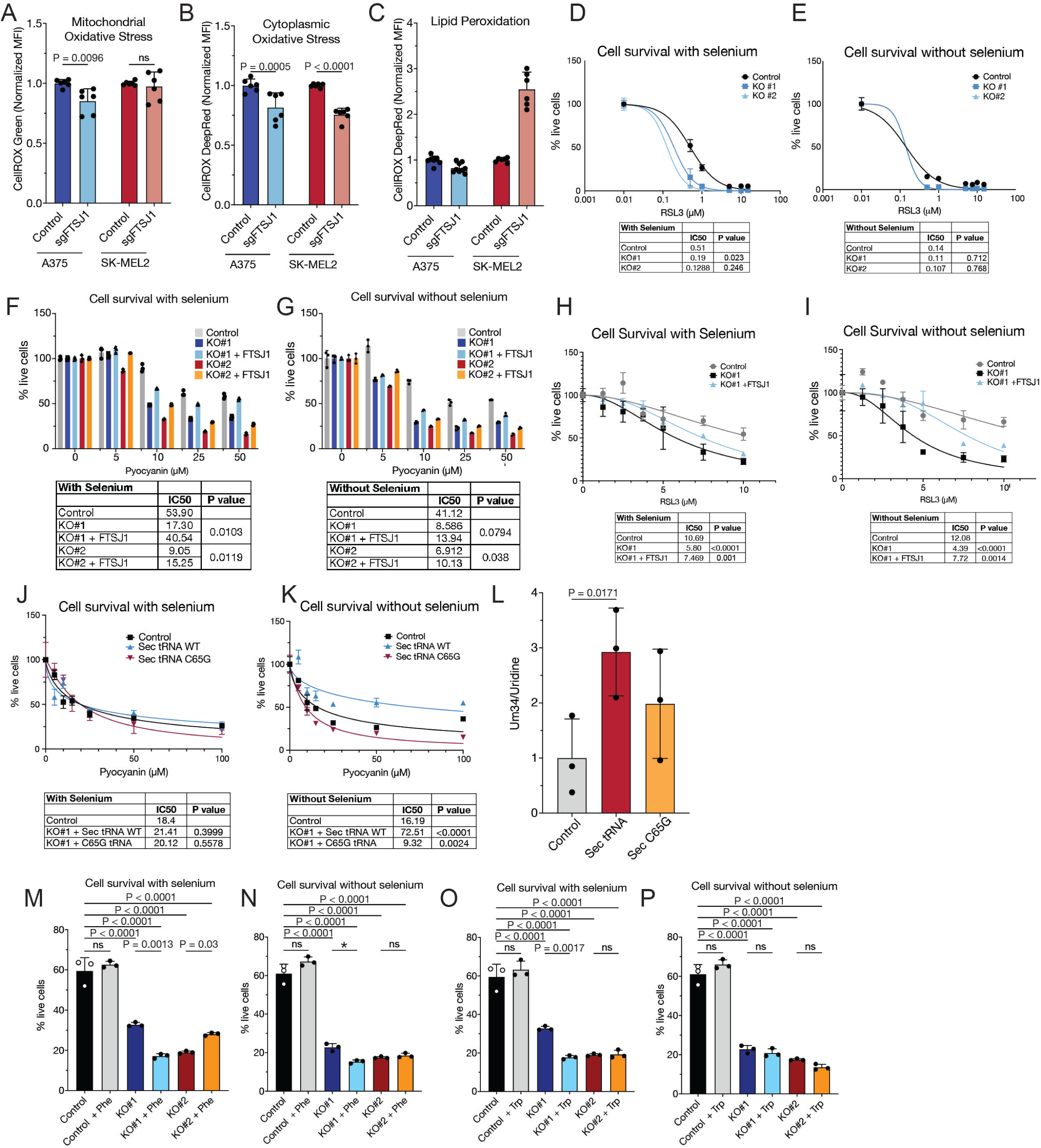

**Extended Figure 8.**
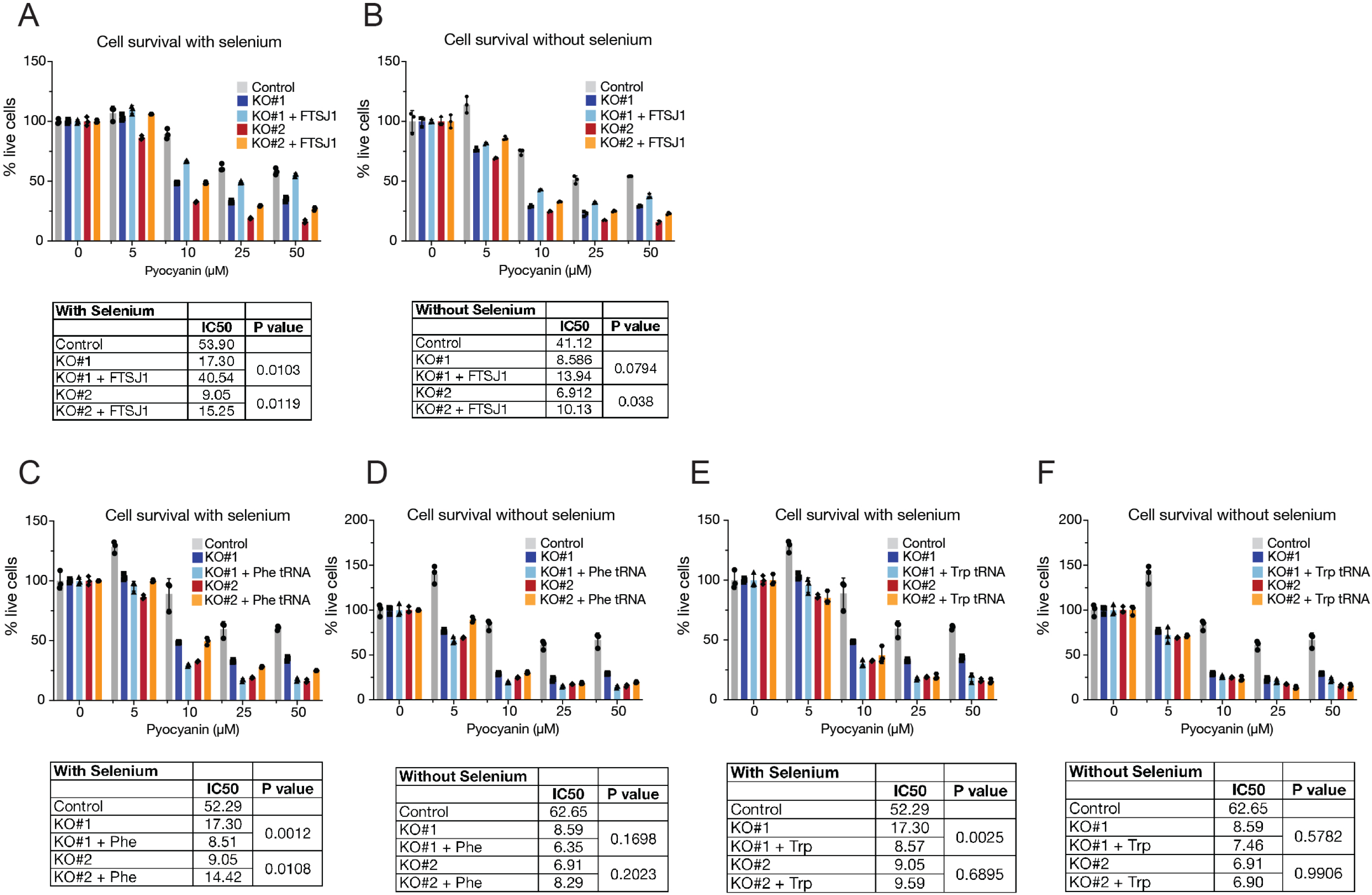

**Extended Figure 9.**
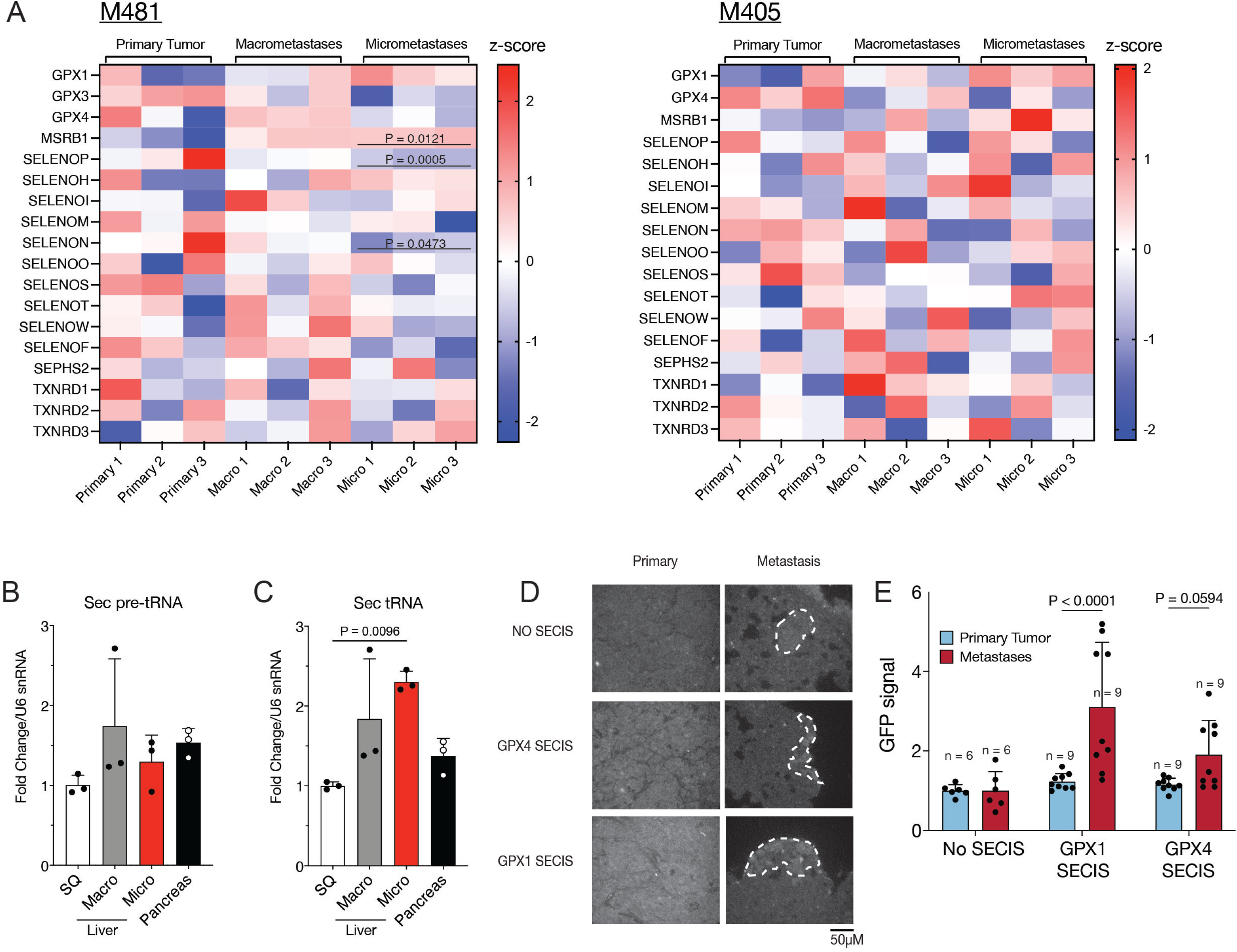

**Extended Figure 10.**
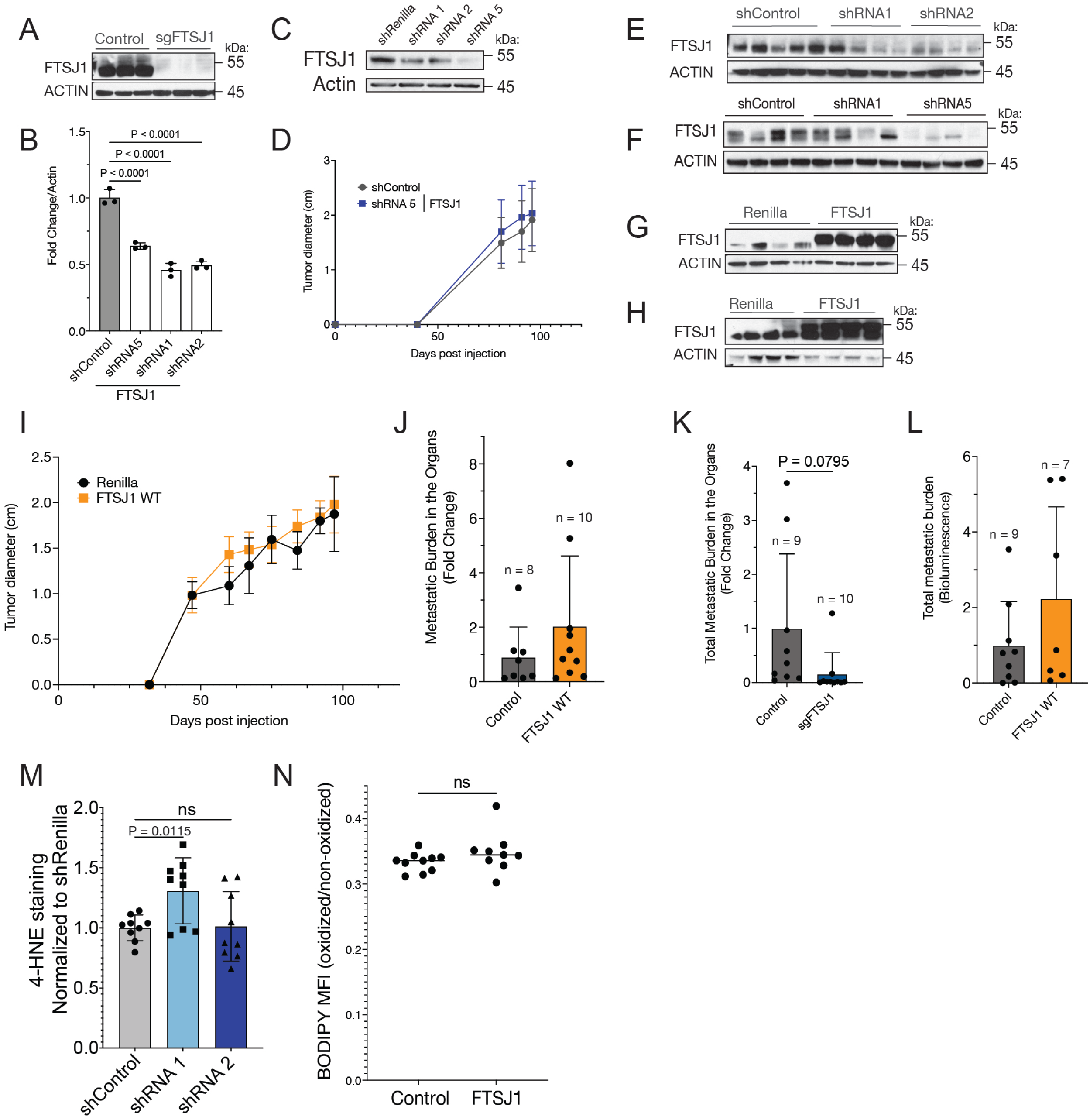

